# Automatic segmentation of medial temporal lobe subregions in multi-scanner, multi-modality magnetic resonance imaging of variable quality

**DOI:** 10.1101/2024.05.21.595190

**Authors:** Yue Li, Long Xie, Pulkit Khandelwal, Laura E. M. Wisse, Christopher A. Brown, Karthik Prabhakaran, M. Dylan Tisdall, Dawn Mechanic-Hamilton, John A. Detre, Sandhitsu R. Das, David A. Wolk, Paul A. Yushkevich

## Abstract

**Objectives:** Volumetry of subregions in the medial temporal lobe (MTL) computed from automatic segmentation in MRI can track neurodegeneration in Alzheimer’s disease. However, poor quality MR images can lead to unreliable segmentation of MTL subregions. Considering that different MRI contrast mechanisms and field strengths (jointly referred to as “modalities” here) offer distinct advantages in imaging different parts of the MTL, we developed a muti-modality segmentation model using both 7 tesla (7T) and 3 tesla (3T) structural MRI to obtain robust segmentation in poor-quality images.

**Method:** MRI modalities including 3T T1-weighted, 3T T2-weighted, 7T T1-weighted and 7T T2-weighted (7T-T2w) of 197 participants were collected from a longitudinal aging study at the Penn Alzheimer’s Disease Research Center. Among them, 7T-T2w was used as the primary modality, and all other modalities were rigidly registered to the 7T-T2w. A model derived from nnU-Net took these registered modalities as input and outputted subregion segmentation in 7T-T2w space. 7T-T2w images most of which had high quality from 25 selected training participants were manually segmented to train the multi-modality model. Modality augmentation, which randomly replaced certain modalities with Gaussian noise, was applied during training to guide the model to extract information from all modalities.

**Results:** The multi-modality model delivered good performance regardless of 7T-T2w quality, while the single-modality model under-segmented subregions in poor-quality images. The multi-modality model generally demonstrated stronger discrimination of A+MCI versus A-CU. Intra-class correlation and Bland-Altman plots demonstrate that the multi-modality model had higher longitudinal segmentation consistency in all subregions while the single-modality model had low consistency in poor-quality images.

**Conclusion:** The multi-modality MRI segmentation model provides an improved biomarker for neurodegeneration in the MTL that is robust to image quality. It also provides a framework for other studies which may benefit from multimodal imaging.

## 1. Introduction

Different MRI contrast mechanisms and field strengths (jointly referred to as “modalities”) offer distinct advantages in imaging different parts of the brain. Multi-modality analysis has been widely used in many brain disease-related tasks, such as brain tumor segmentation (Menze et al., 2015), ischemic stroke lesion segmentation (Lin and Liebeskind, 2016), brain tissue segmentation (Išgum et al., 2015) as well as subregion segmentation in medial temporal lobe (MTL) (Xie et al., 2023).

The MTL, which includes the hippocampus and parahippocampal gyrus, is a frequent focus of research given its role in episodic memory, healthy aging (Maillet and Rajah, 2013; Raz et al., 2004) and brain disorders including neurodegenerative diseases, such as Alzheimer’s disease (AD), (Barkhof et al., 2007; Burton et al., 2009; Korf et al., 2004). The MTL is the earliest cortical region affected by tau proteinopathy, a hallmark pathology of AD (Bilgel, n.d.; Braak and Braak, 1995, 1991; Nelson et al., 2012). Its morphometric abnormalities are not only visible in magnetic resonance imaging (MRI) acquired from patients with AD, but also subtle loss in its subregions can be detected in patients with mild cognitive impairment (MCI) and preclinical AD (Duara et al., 2008; Visser et al., 2002; Xie et al., 2020).

The MTL can be anatomically divided into hippocampal subfields and MTL cortical subregions (called MTL subregions together hereinafter). MRI-based MTL subregional volumetry and morphometry can provide highly sensitive measures for analyzing patterns of neurodegeneration in AD and related disorders (Barkhof et al., 2007; Burton et al., 2009) as well as for mapping of cognition and memory (Squire et al., 2004).

T1-weighted (T1w) 3 tesla (3T) (combined as 3T-T1w) MRI at approximately 1x1x1mm^3^ resolution is the most commonly used MRI contrast for quantifying neurodegeneration. In most large AD neuroimaging studies, such as the Alzheimer’s Disease Neuroimaging Initiative (ADNI) (Petersen et al., 2010) or A4 (Sperling et al., 2020), 3T-T1w modality is the primary structural sequence. However, the appearance of the hippocampus in 3T-T1w MRI lacks sufficient parenchymal contrast, to reliably identify and label the subfields in the hippocampus (Wisse et al., 2021; Yushkevich et al., 2015). On the other hand, a “dedicated” 3T T2-weighted (T2w) (combined as 3T-T2w) sequence offers higher resolution in an oblique coronal plane perpendicular to the hippocampus’s main axis with signal contrast allowing for visualization of subfields (Bonnici et al., 2012; Winterburn et al., 2013). Such dedicated 3T-T2w MRI is being collected in some research protocols, including ADNI, but is less common than 3T-T1w.

The 7 tesla (7T) MRI provides higher resolution and contrast than lower magnetic field scanners for imaging the MTL (Cho et al., 2010; Derix et al., 2014; Prudent et al., 2010; Wisse et al., 2014), and therefore can trace the intricate anatomy of complex regions such as the hippocampal head. It has the potential to identify structural changes in brain disorders with greater accuracy than 3T MRI. Many recent studies utilize the MP2RAGE sequence which combines two volumes acquired at different inversion times, resulting in an image with more inhomogeneity in background but outstanding T1w tissue contrast in the brain (Forstmann et al., 2014). Although 7T-T2w MRI has a high contrast and resolution and can help us observe the subregion structure of MTL clearly (Kollia et al., 2009; Zwanenburg et al., 2011), it is also susceptible to the lateral signal dropout, making the image blurry, especially for the BA36 subregion (Grande et al., 2023). More advantages and disadvantages of these modalities are shown in Table 1.

**Table 1.**
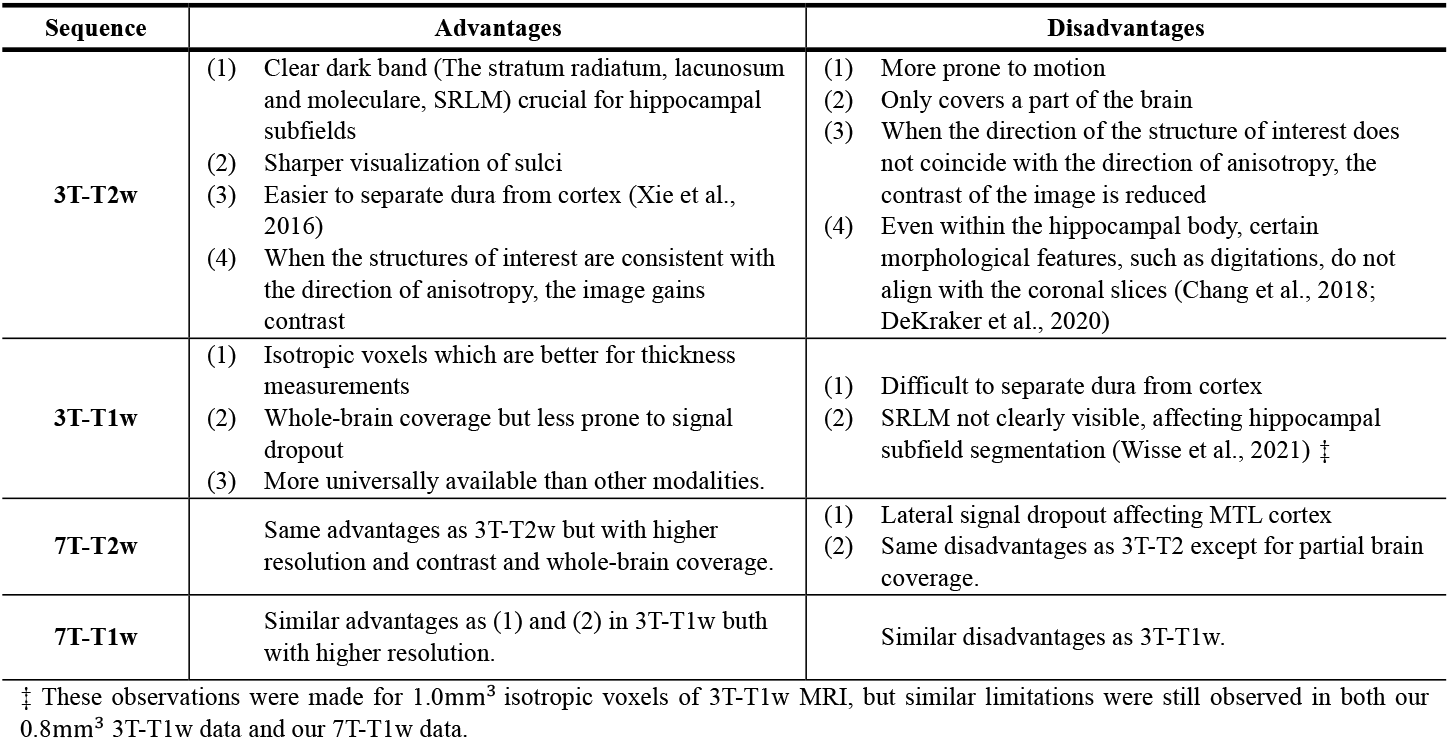
The advantages and disadvantages of different MRI sequences for MTL subregional morphometry.

While 7T MRI offers distinct avantages over 3T MRI in terms of resolution and contrast, image quality of 7T MRI is more variable. In particular, higher magnetic field strength is associated with more severe susceptibility artifacts and therefore some kinds of MRI artifacts are more pronounced at 7T than at 3T. Based on its anatomical location relative to the sinuses, the MTL is particularly vulnerable, especially in the most inferior portion of the MTL with low contrast and low sharpness (see Figure 1). The segmentation difficulties encountered in low-quality images are due to the lack of sufficient anatomical information for segmentation, so that neither experienced experts nor segmentation models can complete accurate segmentation. To overcome this issue, it is necessary to consider additional information outside of 7T-T2w using multiple available modalities.

**Figure 1.**
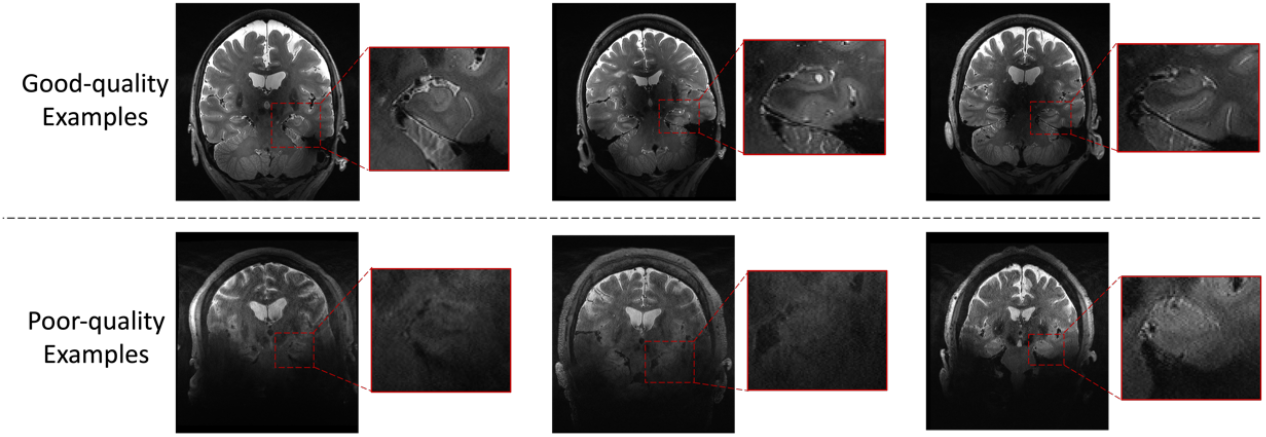
Examples of good image quality and poor image quality in 7T-T2w modality.

Segmenting the MTL subregions manually is a challenging and time-consuming task, which is prone to inter-rater differences and requires a lot of anatomical knowledge (Hickling et al., 2024; Olsen et al., 2017; Yushkevich et al., 2015). Automatic segmentation of the MTL subregions is less laborious and more reproducible. The existing segmentation algorithms for hippocampal subfields or MTL subregions can be categorized into two groups: template/atlas-based methods and deep learning-based methods.

The template/atlas-based methods can be further divided into statistical models such as FreeSurfer (Iglesias et al., 2015; Van Leemput et al., 2009); surface-based models such as SurfPatch (Caldairou et al., 2016) and HippUnfold (DeKraker et al., 2022); and registration-based models such as ASHS (Wisse et al., 2016; Xie et al., 2019; Yushkevich et al., 2015), LASHiS (Shaw et al., 2020), and HIPS (Romero et al., 2017). Deep learning based MTL subregion segmentation models include Deep Label Fusion (Xie et al., 2023), deep HIPS (Manjón et al., 2022), and the weak network fusion method (Poiret et al., 2023). However, most of these existing methods only consider a single MRI modality. While ASHS, SurfPatch and HippUnfold consider both T1 and T2, the T1 modality is used to assist in the registration of T2-weighted MRI and not directly for segmentation step. HIPS uses both T1 and T2, but it was developed in single magnetic field strength and the authors did not verify its performance in 7T MRI, which is more prone than 3T MRI to motion-related and other artifacts.

In this study, we have developed an integrative model that harnesses data from all available MRI modalities, including 3T-T1w, 3T-T2w, 7T-T1w, and 7T-T2w, and we verified that using multi-modality data can enhance segmentation performance, especially when the quality of the primary modality is poor (the primary modality is the modality in the space of which the segmentation is generated; this distinction is necessary because both at 3T and 7T, the T1-weighted and T2-weighted scans typically have different orientation and resolution). Our model is designed to fuse information across the available modalities and to retrieve complementary data when one or more modalities are compromised or incomplete. Drawing upon the robust performance of nnU-Net in the context of hippocampal subfield segmentation as demonstrated in (DeKraker et al., 2022; Xie et al., 2023), coupled with the prevalent adoption of deep learning techniques in medical imaging analysis, we have chosen to implement nnU-Net as the baseline model on which to evaluate the impact of including multiple modalities on segmentation performance.

Our study is carried out in a large dataset (n=197) of 3T and 7T MRI in the same set of research study participants at the University of Pennsylvania Alzheimer’s Disease Research Center (ADRC). The input for the model consists of multi-modality images co-registered and resliced to the space of the 7T-T2w image. A modality augmentation scheme is proposed to force the model to extract useful features from all modalities. The nnU-Net is used as the segmentation backbone. The model’s performance is first evaluated through cross-validation on the available set of manually-labeled atlases. However, recognizing that evaluation in this high-quality image dataset is likely not representative of real-world performance on individuals whose 7T-T2w scans are of poorer quality, we perform three additional indirect evaluations, including (1) volume comparison, (2) longitudinal comparison and (3) a task-based evaluation comparing the ability of different models to distinguish Amyloid-cognitively unimpaired (A-CU) and Amyloid+ MCI (A+MCI) individuals. Together, these indirect evaluations demonstrate that the proposed multi-modality segmentation model outperforms the baseline single-modality model that only considers the 7T-T2w modality, especially for poor quality 7T-T2w images.

This study is an application of using multi-modality images to reduce the impact of poor quality images on segmentation performance. The evaluation assessed the model’s performance on a training set with high-quality images and a test set containing many low-quality images, a situation that is likely common in various medical imaging domains. The challenges addressed in this paper are relevant to other medical image analysis applications in which multi-modality data is available, including other MRI contrasts such as diffusion tensor imaging (DTI) and susceptibility weighted imaging (SWI). The code of this study is available at https://github.com/liyue3780/mmseg.

## 2. Materials and Methods

The flowchart of the training process of the segmentation model is shown in Figure 2. It includes whole-brain registration, ROI determination, local registration, modality augmentation, and nnU-Net training. In the model inference, the modality augmentation will be skipped and the model will use full modality as input. Details of each step are explained below.

**Figure 2.**
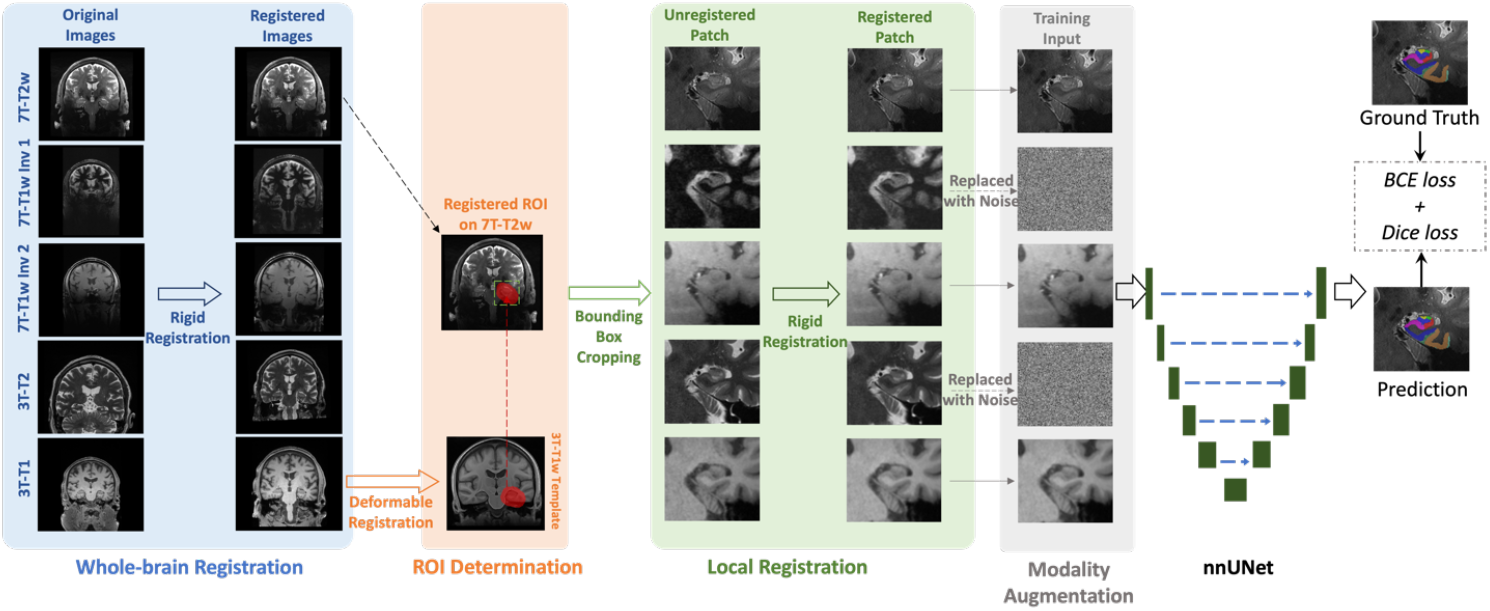
The flowchart of the training process of the segmentation model. All modalities are rigidly registered to 7T-T2w in the whole-brain registration step. Then, the MTL ROI (red region in ROI determination step) is mapped from 3T-T1w template to 7T-T2w space by deformable registration. MTL patches are cropped along the bounding box of the ROIs in all registered modalities and local registration step refines the misalignment between 7T-T2w and other modalities specifically on MTL. Before feeding into the nnU-Net segmentation model, one to four modalities (two modalities in this figure as examples) are randomly replaced by noise in modality augmentation step. Finally, the nnU-Net is trained using combined BCE and Dice as loss function

### 2.1 Dataset

#### 2.1.1 Participants and MRI protocol

MRI scans were acquired from the ADRC at the University of Pennsylvania. Both 7T and 3T MRI were considered in this study. The 7T protocol included both T1w (MP2RAGE) and T2w (TSE) scans. The 7T-T1w scan contained two inversion contrasts with different T1 weightings, INV1 and INV2, which had a resolution of 0.69 × 0.69 × 0.69 mm^3^. The 7T-T2w scan covered the whole brain and had in-plane resolution of 0.42 × 0.42 mm^2^ and slice thickness of 1 mm.

The 3T protocol included a ‘routine’ whole-brain T1w (MPRAGE) scan and a ‘dedicated’ T2w (TSE) scan with partial brain coverage. The resolution of the 3T-T1w scan was 0.8 × 0.8 × 0.8mm^3^. The in-plane resolution and slice thickness of the 3T-T2w scan were 0.4 × 0.4 mm^2^ and 1.2 mm, respectively, with the slice orientation approximately orthogonal to the main axis of the hippocampi.

The inclusion and exclusion criteria used in this study are shown in Supplementary Section S1. After excluding the participants that were not diagnosed as cognitively unimpaired (CU) or MCI, had inconsistent diagnosis in 3T and 7T sessions, had modality missing, and had incorrect registration (described in Section 2.2), we finally obtained 25 training participants and 172 test participants.

The most recent diagnosis result and amyloid PET to the date of 7T scan of each participant were collected. Of these participants, 40 were diagnosed with MCI, while the remaining 157 were cognitively unimpaired (CU) individuals, 122 of them had Amyloid PET with 40 diagnosed as positive and 82 negative scans based on visual read. Table 2 shows specific cohort characteristics in this study.

**Table 2.** Characteristics of the cohort in this study. (Abbreviations: PET=positron emission tomography, ROI=region of interest, CU= cognitively unimpaired, MCI= mild cognitive impairment)

|  | Training Set |  | Test Set |  |
| --- | --- | --- | --- | --- |
| Number of participants | 25 |  | 172 |  |
| Age at the time of 7T scan | 59 – 97<br>(72.2 ± 7.5) |  | 23 – 94<br>(63.0 ± 16.7) |  |
| Sex (female/male) | 16/9 |  | 103/69 |  |
| High-quality 7T-T2w ROIs † | 47 |  | 231 |  |
| Low-quality 7T-T2w ROIs | 3 |  | 113 |  |
| CU participants | 15 | (Amyloid–: 9)<br>(Amyloid+: 3) | 142 | (Amyloid–: 65)<br>(Amyloid+: 15) |
| MCI participants | 10 | (Amyloid–: 2)<br>(Amyloid+: 4) | 30 | (Amyloid–: 6)<br>(Amyloid+: 18) |
| Amyloid PET | 18 |  | 104 |  |
| Longitudinal scan | - |  | 29 |  |

† Because each participant has two ROIs, left and right, the total number of ROIs is twice the total number of participants. The quality assessment is described in section 2.1.3

Among 172 test participants, some of them had more than one 7T scan. Using the inclusion and exclusion criteria described in Supplementary S1, longitudinal scans of 29 participants were collected.

The distributions of scanning date interval between 7T and 3T MRI of 172 test participants and 25 training set, respectively, scanning interval between two longitudinal 7T scans in 29 participants with longitudinal data are shown in Supplementary Figure S2.

#### 2.1.2 Gold standard Annotation for Training Set

A total of 25 participants were previously selected as the ASHS “atlas set” and underwent manual segmentation (Xie et al., 2023). As shown in Table 2, most ROIs in atlas set had high quality (the quality assessment process is described in Section 2.1.3). This subset also serves as the training set for the proposed segmentation model.

The “Penn Aging Brain Cohort (ABC)” protocol, originally described in (Berron et al., 2017), and modified in (Xie et al., 2023) was used to segment the hippocampal subfields and cortical MTL subregions manually on both left and right sides in 7T-T2w scans of training set. The specific subregions are introduced in Table 3 and shown in Figure 3.

**Table 3.** Summary of subregions manually labelled according to ABC protocol (Berron et al., 2017)

| Category | Abbr. | Name | Anatomical Explanation |
| --- | --- | --- | --- |
| Hippocampal Subfields | CA1-3 | Cornu ammonis 1-3 | The CA region is located laterally from the subiculum and wraps around the DG. It consists of several subdivisions named CA1, CA2 and CA3. A distinction between these CA subfields can be made based on the size, shape and density of the pyramidal cells (Henri Duvernoy, 2005). CA subfields have three layers: 1) the alveus, a white matter layer, 2) a pyramidal layer, and 3) the stratum radiatum lacunosum moleculare, another white matter layer (Henri Duvernoy, 2005). The white matter layers are visible on high resolution T2w MRI and help visualize the inner structure of the hippocampus. |
|  | DG | Dentate gyrus | The DG is mostly encapsulated by the CA regions and to some extent the subiculum. The DG consists of a molecular layer, a granular cell layer and a polymorphic layer. The DG also includes the subfield hilus. The DG can be distinguished on T2w MRI from surrounding CA and subiculum regions because the white matter layers of these regions. |
|  | SUB | Subiculum | The subiculum is a pivotal structure located between the hippocampus proper (CA and DG), and entorhinal cortex and other cortices (O'Mara et al., 2001). It consists of several layers: 1) the alveus, a white matter, 2) a pyramidal cell layer, and 3) a polymorphic layer (Ding, 2013). As for CA, the white matter layers are visible on high resolution T2w MRI. In this segmentation protocol the subiculum label also includes presubiculum and parasubiculum, which have a more complex laminar organization (Insausti and Amaral, 2012). |
|  | Tail | Tail | The tail is the posterior part of the hippocampus which consists of the same subfields as the anterior hippocampus, including CA1-3, DG and the subiculum. It is not partitioned into subfields in our segmentation protocol because of its complex anatomy and curvature of the hippocampal axis relative to the imaging plane. |
| Extra-hippocampal Cortical Subregions | ERC | Entorhinal cortex | The ERC is located anteriorly on the crown of the parahippocampal gyrus, adjacent, inferiomedially, to the subiculum. It consists of six layers with a distinct cell-free layer IV, the lamina dissecans (Insausti and Amaral, 2012). The ERC is a multimodal limbic association area in MTL. It receives the projections from other multimodal areas, secondary association areas and limbic structures. |
|  | BA35 | Brodmann area 35 | BA35 and BA36 constitute the PRC, which is a region in anterior MTL. The PRC is located laterally from the ERC, but also wraps around the anterior and posterior end of the ERC. BA35 is mostly located in the collateral sulcus, whereas BA36, laterally from BA35, extends often into the fusiform gyrus (Ding and Van Hoesen, 2010). BA35 and BA36 have 6 layers but no lamina dissecans (Wuestefeld et al., 2024). |
|  | BA36 | Brodmann area 36 |  |
|  | PHC | Parahippocampal cortex | The PHC makes up the posterior part of the parahippocampal gyrus and therefore borders the posterior subiculum inferiomedially. The anterior border of the PHC consists of the ERC and PRC. PHC also contains six layers. |
| Non-gray-matter Labels | HS | Hippocampal sulcus | - |
|  | CS | Collateral sulcus | - |
|  | Cysts | Cysts | - |
|  | MISC | Miscellaneous | - |

**Figure 3.**
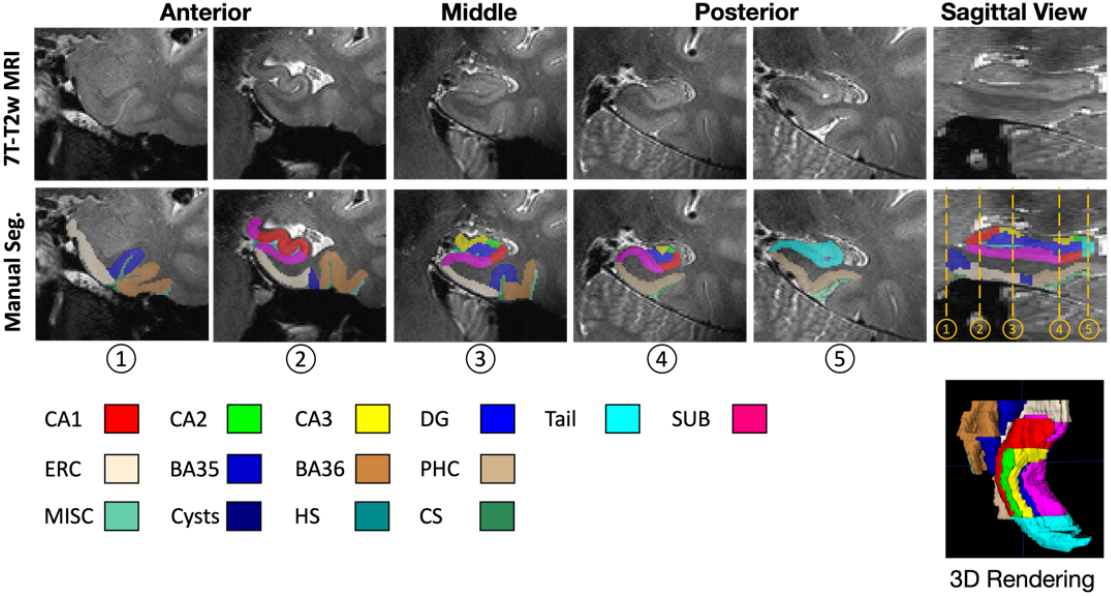
Examples of gold standard annotation in the training set.

The training atlas including left and right MTL ROIs and manual segmentations has been shared under the Creative Commons Zero license (download at https://doi.org/10.5061/dryad.0zpc8676p)

The remaining 172 participants without gold standard (manual segmentation) were used as the test set to evaluate the model. The test set was not selected according to image quality, and therefore contained participants with poor 7T-T2w image quality, which is common in real-world 7T-T2w dataset.

#### 2.1.3 Quality Assessment

To evaluate segmentation performance across different image qualities, the image quality of 7T-T2w modality for both training and test sets was assessed and labeled by author YL. The participants in the training and test sets were mixed together and randomly shuffled for assessment. The assessment process was blinded to the rater. The specific steps are as follows:

- The MTL region of each participant was cropped from the 7T-T2w MRI.
- The participants were inspected in random order.
- The rater navigated through the ROIs in 3D in ITK-SNAP (Yushkevich et al., 2006) and rated them on a scale of 1 to 9 based on the visibility and sharpness of hippocampus and cortical regions, with higher numbers indicating better image quality. When assessing, rater used the number 5 as a threshold, and all images considered to be of high quality would have a score greater than or equal to 5.
- The assessments were conducted three times and the average rating of each ROI was calculated to minimize bias.
- After rounding, the rating will be used as the final rating for each participant.

### 2.2 Pre-processing

#### 2.2.1 Whole-brain Rigid Registration

The different modalities had different resolutions (as displayed in Figure 2), voxel spacing and slice thicknesses. All modalities’ images were registered to primary modality (7T-T2w) space rigidly using the registration tool *greedy* (Venet et al., 2021) in each participant.

This process in each participant can be divided into three steps.

- ***7T-T1w to 7T-T2w*:** The 7T-T1w INV1 and 7T-T1w INV2 had same resolution and voxel spacing. They were registered together to 7T-T2w with normalized mutual information (NMI) as image similarity metric, center alignment by translation as initial registration, and 100, 50 and 10 as number of iterations at coarsest level (4x), intermediate (2x) and full (1x) resolution, respectively.
- ***3T-T1w to 7T-T2w*:** The 3T-T1w modality was registered to 7T-T2w rigidly with NMI as image similarity metric, center alignment by translation as initial registration, and 100, 50 and 10 as number of iterations at coarsest level (4x), intermediate (2x) and full (1x) resolution, respectively.
- ***3T-T2w to 7T-T2w:*** The 3T-T2w image was “dedicated” scan with partial brain coverage that targets the hippocampal region specifically. Performing registration from 3T-T2w to 7T-T2w directly could result in misalignment because the uncovered region (zero-value voxels) were involved in the calculation of registration similarity. Since 3T-T1w and 3T-T2w were acquired in the same scan, they were roughly aligned. The rigid registration matrix calculated from 3T-T1w to 7T-T2w was used to map 3T-T2w image to 7T-T2w space.

Registration accuracy was visually checked. The examples of successful and failed registrations are shown in Supplementary Figure S3.

#### 2.2.2 Region of Interest Determination

The current study focuses on MTL subregions. Thus, the regions surrounding the left and right MTL were selected as regions of interest (ROI), respectively. An unbiased population template (Joshi et al., 2004) constructed using 29 3T-T1w MRI scans which were from a different population with this study, and template-space ROIs for the left and right MTL, generated as part of the 3T-T1w ASHS package (Xie et al., 2019), were used in this process.

The ROI determination for each participant includes the following steps.

- First, the neck was trimmed in the 3T-T1w image.
- Then, the template was registered to the neck-trimmed 3T-T1w image using deformable registration (using *greedy* (Venet et al., 2021)) with the similarity metric of normalized cross-correlation (NCC) calculated with the neighborhoods of 5x5x5 voxels. The deformation field was saved and the left and right ROIs in template were mapped to the 3T-T1w image using the deformation field.
- Since the 3T-T1w image had been registered to 7T-T2w space in step 2.2.1, these ROIs were further mapped to the 7T-T2w space using registration matrix obtained in 2.2.1.
- Left and right MTL patches along the bounding box of the registered ROIs were cropped from 7T-T2w image and other modalities’ images in 7T-T2w space according to the ROIs in 7T-T2w.

Registration accuracy was visually checked. The examples of successful and failed registrations are shown in Supplementary Figure S4.

The ROI sizes varied among 197 participants. On average, the ROIs measured 72.58mm in the anterior-posterior direction with a range from 62mm to 87mm, 60mm in the inferior-superior direction with a range from 43.68mm to 61.07mm, and 55.01mm in the left-right direction with a range from 48.35mm to 61.92mm.

#### 2.2.3 Local Rigid Registration

The whole-brain affine registration was not sufficient to align the local MTL region well due to modality-related non-linear deformations, e.g., MRI gradient distortion. This motived the need for further local registration. Therefore, a more reliable rigid registration that focused only on left and right ROIs, respectively, was performed by *greedy* (Venet et al., 2021).

The local registration can be divided into three steps.

- ***7T-T1w to 7T-T2w***: 7T-T1w inv1 had similar intensity distribution to 7T-T2w, so the NCC calculated in neighborhoods of 5x5x5 was chosen as similarity metric for the rigid registration from 7T-T1w INV1 patch to 7T-T2w patch. Then, the 7T-T1w INV2 patch was mapped to 7T-T2w using the same registration matrix as 7T-T1w INV1.
- ***3T-T1w to 7T-T2w***: the 3T-T1w patch was rigidly registered to the 7T-T2w with NMI as similarity metric, identical alignment as initialization, and 100 and 50 as number of iterations at the coarsest level and intermediate resolution, respectively.
- ***3T-T2w to 7T-T2w***: 3T-T2w image had similary intensity distribution as 7T-T2w. 3T-T2w patch was rigidly registered to 7T-T2w with weighted-NCC as similarity metric, identical alignment as initialization, and 100 and 50 as number of iterations at the coarsest level and intermediate resolution, respectively.

This resulted in two ROIs for each participant with each ROI containing five well-registered modalities. Registration accuracy was visually checked. The examples of successful and failed registrations are shown in Supplementary Figure S5.

### 2.3 Segmentation Model

The current study focuses on leveraging multiple MRI field strengths and modalities in MTL subregion segmentation rather than on developing a novel segmentation backbone. We conducted experiments using the extensively validated nnU-Net framework (Isensee et al., 2021) as the segmentation model. Specifically, the 3D full-resolution version of nnU-Net was selected.

The input of the model was a five-channel 3D input (as per the order, the five channels were 7T-T2w, 7T-T1w inv1, 7T-T1w inv2, 3T-T2w, and 3T-T1w) which was formed by stacking all modalities in the same ROI with 7T-T2w. All ROIs on the right were flipped to the left side before feeding into the segmentation model.

Due to the high quality of the 7T-T2w modality in the training set and the fact that manual segmentation was based on the 7T-T2w, we would expect the nnU-Net model to primarily rely on the 7T-T2w to minimize its loss during training and largely ignore information from other modalities. However, when applied to the larger testing set, where the average quality of the 7T-T2w is lower, we hypothesize that a model that primarily relies on 7T-T2w would underperform compared to a model that effectively synthesizes information from all available modalities. To encourage nnU-Net to use all available modalities during training, we incorporate an additional augmentation scheme, called *modality augmentation (ModAug)*, which we employ together with the standard data augmentation schemes in nnU-Net.

As shown in Figure S6 in supplementary material, in each iteration of training process, there was a 50% chance that input data was fed into an augmentation branch. Otherwise, all five modalities were fed into the segmentation model. In the augmentation branch, there was an equal chance that data were processed by each of four different sub-branches, where one, two, three or four modalities in the input data would be replaced by random noise. This step was designed to force the model to extract critical information from all modalities, rather than focusing only on the information associated with the primary modality. In addition to random noise, all-zero images and images from other participants were also used in ModAug. The details and performance of these are shown in supplementary material S10. Due to their similar performance, the model used in this paper implemented ModAug using random noise.

Combined Dice and binary cross-entropy (BCE) loss function was used to supervise the training process using deep-supervision scheme. The model was trained in a five-fold cross-validation setting. In each fold, the training process was terminated after 400 epochs, where the validation loss of the validation set plateaued to constant. All other hyper-parameters remained at their default settings. When performing inference on the test set without manual segmentations (n=172), the five per-fold models were ensembled to generate the final prediction (Isensee et al., 2021).

### 2.4 Model Evaluation

#### 2.4.1 Baseline Model

To verify the advantage of using multiple modalities, another nnU-Net model which only used the primary modality (7T-T2w modality) as input was trained as baseline model. It is referred to as the single-modality model.

#### 2.4.2 Cross-validation in the Annotated Training Set

As a metric to measure the consistency between predicted segmentation and gold standard, Dice similarity coefficient (Dice, 1945) was used to evaluate segmentation accuracy in five-fold cross-validation in the training process. For each subregion, the average of Dice scores from all validation participants in five folds were calculated.

#### 2.4.3 Validation in the Test Set

Due to the variable quality of the 7T-T2w images and the significant expert effort it would entail, manual segmentations of the test set were not conducted. As a result, direct quantitative evaluation, such as Dice score, could not be calculated. Instead, one qualitative evaluation, along with downstream evaluation methods, was used to compare the segmentation performance of the proposed model and the baseline model.

- ***Qualitative evaluation***. Screenshots of segmentation results of single-modality model and multi-modality model were generated, respectively, using standard quality control pipeline in ASHS (Yushkevich et al., 2015). For each side of the MTL ROI, nine equally spaced coronal slices and two sagittal slices of the 7T-T2w images as well as corresponding segmentation results were displayed in the screenshot. All screenshots for these two models were mixed and randomly shuffled. A blind rating was conducted by author Y.L. in the random sequence. For each segmentation, the rater assessed it visually and classified it into three categories: good, bad, and uncertain (in between good and bad segmentation). Confusion matrix was calculated to show the segmentation quality comparison between these two models.
- ***Longitudinal consistency***. A good segmentation model should perform well regardless image quality. When presented with two longitudinal scans for the same participant in our study, a good model should provide largely consistent MTL subregion volume measurements, considering that the loss of hippocampal MTL volume in healthy aging and MCI in the range of 0.2-2.55% a year (Fjell et al., 2009; Jack et al., 2000; Kurth et al., 2017). Intraclass correlation coefficients (ICC) was used to evaluate longitudinal consistency for each model in the subset of 29 participants who had longitudinal 3T and 7T MRI.
- ***Ability to discriminate between CU and MCI***. Previous studies showed that regional measurements such as cortical thickness and subfield volumes had the ability to discriminate between A+MCI participants and A-CU participants (Whitwell et al., 2007; Wolf et al., 2004). Correct computation of measurements relies on accurate segmentation of subregions, so the ability to separate A-CU and A+MCI was used as a metric for evaluating segmentation models. Specifically, a general linear model (GLM) was fitted in each subfield or subregion. Each GLM used diagnostic group (A+MCI and A-CU) as the independent variable. For the subfields of the hippocampus, the GLM’s dependent variable was subfield volume and the GLM included age and intracranial volume (ICV) as nuissance covariants; for the cortical subregions, the GLM’s dependent variable was median thickness calculated by computing the pruned Voronoi skeleton (Ogniewicz and Kübler, 1995) of the MTL cortex and integrating the radius field over each MTL subregion, as implemented in the software CM-Rep (Pouch et al., 2015), and only age was included as the nuissance covariant.
- ***Generalization performance***. The ability of the proposed pipeline generalized to a new dataset (Chu et al., 2025) from different scanners was evaluated.

To ensure comparability across participants in volume-based evaluation, each hippocampal subfield volume was adjusted to account for the inclusion of the hippocampal tail, based on the ratio of the total hippocampal volume to the volume excluding the tail. Additionally, the volumes of the MTL cortical subregions were normalized using the physical length along the anterior-posterior direction, as suggested in (Yushkevich et al., 2015).

## 3. Results

### 3.1 Quality Assessment

Image quality was first assessed so that we could evaluate the performance of the models in images of different quality. Left and right ROIs of all participants in the whole dataset were assessed. Figure 4 shows the distribution of image quality in the training set and the test set. The rating greater or equal to 5 represents the good quality and less than 5 represents the poor quality.

**Figure 4.**
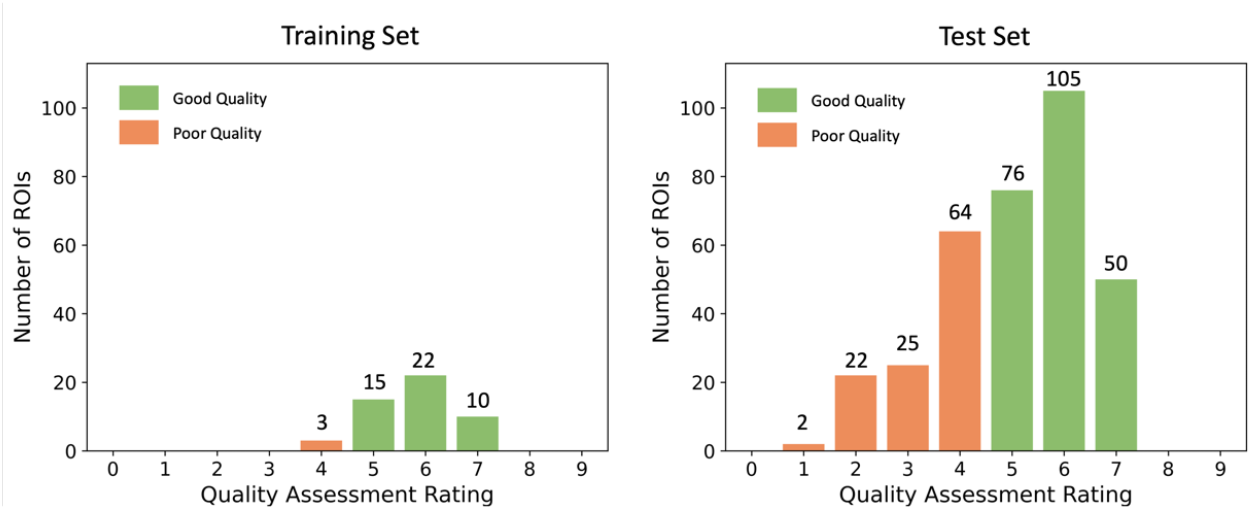
The distribution of image quality in the training and test set respectively.

The average quality of the training set was 5.79, while the average quality of the test set was 5.04. The majority of the ROIs in the training set had good image quality. Only three ROIs in the training set (6%) had quality rating 4 and none were rated below 4. As expected, the test set had a substantially larger proportion of poor quality ROIs (32.85% with rating lower than 5). However, the distribution of the quality ratings in the [5,9] range was similar for the test set and the training set (the ratio of samples rated 5, 6 and 7 in both sets was approximately 3:4:2).

Therefore, cross-validation in the training set does not fully demonstrate a model’s performance in real-world data. We still need to evaluate the model’s segmentation ability in low-quality images in the test set.

### 3.2 Cross-validation in the Training Set

Dice coefficient between automated and manual segmentation was calculated for each subregion in the cross-validation experiments. Table 4 compares the average Dice coefficient for each subregion among the single-modality models (7T-T2w as input, 7T-T1w as input, 3T-T2w as input, and 3T-T1w as input), and multi-modality models (with and without ModAug in training process). For these single-modality models, the segmentation was performed in space of 7T-T2w. All other modalities were registered to 7T-T2w before we trained and test these models.

**Table 4.** Dice score (average Dice ± standard deviation) comparison of single-modality and multi-modality models in five-fold cross-validation (Bold numbers represent the highest values among all models)

| Subregions | Single-modality Models |  |  |  | Multi-modality Models |  |
| --- | --- | --- | --- | --- | --- | --- |
|  | 7T-T2 | 7T-T1 | 3T-T2 | 3T-T1 | without ModAug | with ModAug |
| CA1 | 0.814 | 0.765 *** | 0.767 *** | 0.758 *** | **0.817** | **0.817** |
| CA2 | 0.735 | 0.686 *** | 0.682 *** | 0.675 *** | **0.747** | 0.741 |
| CA3 | 0.705 ** | 0.685 *** | 0.674 *** | 0.679 *** | **0.728** | 0.724 |
| DG | 0.870 ** | 0.813 *** | 0.814 *** | 0.807 *** | **0.871 \*\*** | 0.866 |
| SUB | 0.861 | 0.822 *** | 0.818 *** | 0.816 *** | **0.862** | **0.862** |
| Tail | 0.845 *** | 0.833 *** | 0.823 *** | 0.827 *** | 0.859 | **0.861** |
| ERC | 0.846 *** | 0.810 *** | 0.802 *** | 0.809 *** | 0.845 *** | **0.855** |
| BA35 | 0.730 *** | 0.709 *** | 0.691 *** | 0.704 *** | 0.727 *** | **0.751** |
| BA36 | 0.812 *** | 0.781 *** | 0.764 *** | 0.778 *** | 0.802 *** | **0.825** |
| PHC | 0.815 | 0.792 *** | 0.778 *** | 0.789 *** | 0.818 | **0.822** |

\*\* $p < 0.01$ ; \*\*\* $p < 0.001$ in paired t-test comparison between corresponding model and multi-modality model with ModAug

Among all the single-modality models, the 7T-T2 model had the best cross-validation performance. Its Dice scores were much higher than other three single-modality models, while the other three had very comparable performance.

For each subregion, the highest Dice value was obtained from multi-modality model, either with ModAug or without ModAug. It shows the advantage of involving other modalities in a segmentation model, which indeed provides more useful information than any single modality alone.

The comparison between two mutli-modality models with and without ModAug shows the model without ModAug performed better in hippocampus subfields and the model with ModAug performed better in extrahippocampal subregions. The p-value shows the segmentation on hippocampus was comparable between these two models in all subfields except for DG, while the model with ModAug significantly performed better in ERC, BA35 and BA36. This indicates the ModAug encouraged the model to learn multi-modality information better.

### 3.3 Qualitative evaluation

Table 5 shows the confusion matrix of qualitative evaluation for single-modality model and multi-modality model with ModAug. For the single-modality model, 36 segmentations were rated bad, and 70 uncertain, but for the multi-modality model only one was rated bad and 43 uncertain. The multi-modality model rescued 31 of 36 “bad” segmentations and 42 of 70 “uncertain” segmentations.

**Table 5.** Confusion matrix of segmentation quality between single-modality and multi-modality models.

| Single-modality \ Multi-modality | Good | Uncertain | Bad | Total |
| --- | --- | --- | --- | --- |
| Good | 227 | 11 | 0 | 238 |
| Uncertain | 42 | 28 | 0 | 70 |
| Bad | 31 | 4 | 1 | 36 |
| Total | 300 | 43 | 1 | 344 |

To show how these two models varied with the quality of the image, seven examples with different image qualities in the test set and two examples with high quality in the training set were randomly selected. Figure 5 shows these examples. When the image quality was poor, the single-modality model exhibited under-segmentation while the multi-modality model could still segment all subregions resonably. As the image quality improved, the under-segmentation of single-modality model became less and its segmentation results approached the multi-modality model.

**Figure 5.**
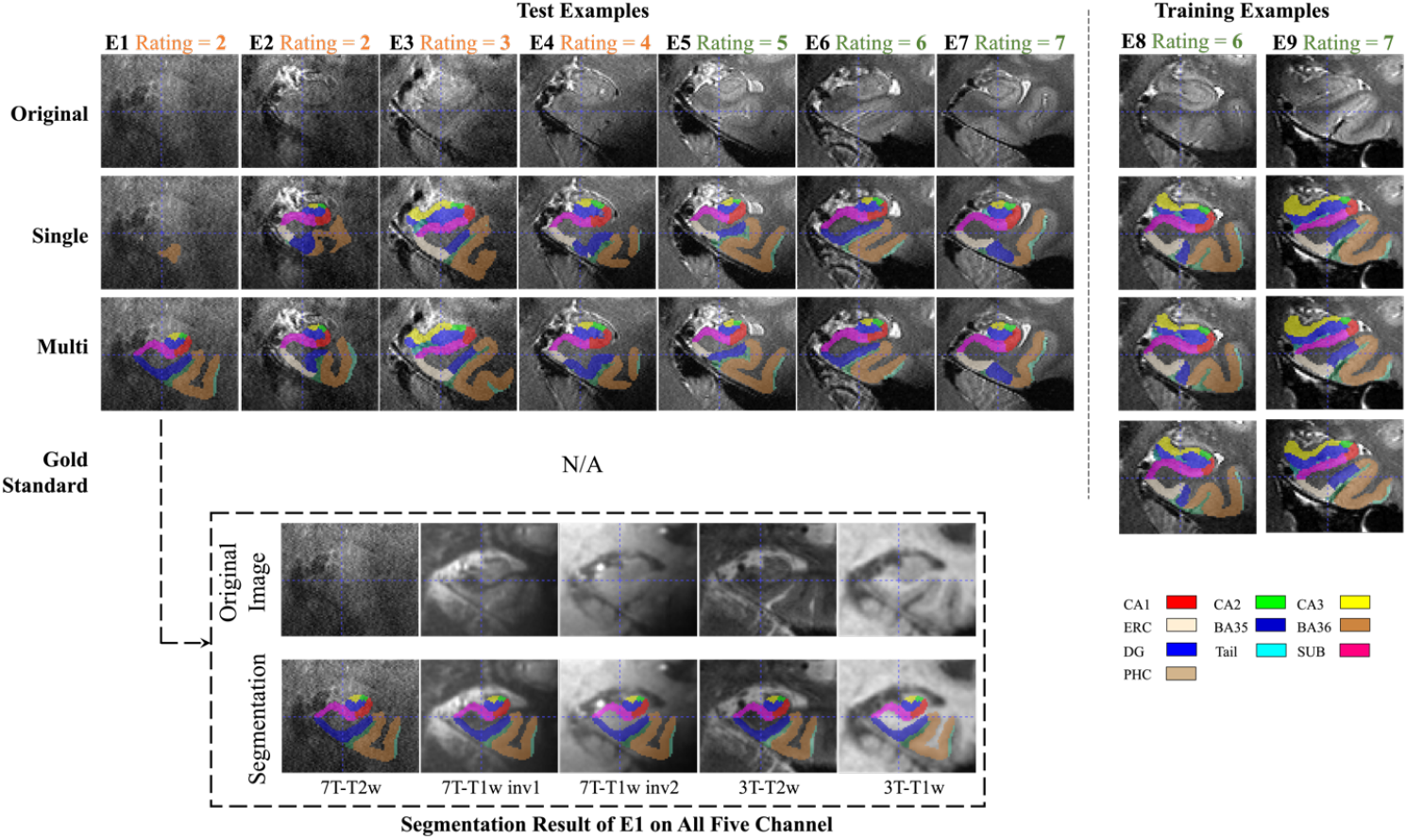
Subregion segmentation in examples with imaging quality from low to high. The segmentation in training examples are obtained by cross-validation. The higher the image quality, the more consistent between single-modality and multi-modality models. For the first example (E1), other modalities are shown in dashed box and the segmentation by multi-modality model aligns well in these modalities. (CA = cornu ammonis; DG = dentate gyrus; SUB = subiculum; ERC = entorhinal cortex; BA = Brodmann area; CS = collateral sulcus; HS = hippo sulcus; MISC = miscellaneous label)

Figure S12 in the supplementary materials shows a comparison of subregion volumes, while supplementary Table S2 presents a comparison of segmentation quality and image quality. Both provide further evidence of the robustness of the multi-modality model across different image qualities.

### 3.4 Longitudinal Comparison

For the 29 test participants with longitudinal MRI scans, the ICC of volumes obtained from different timepoints of each subregion were computed. Additionally, the Bland-Altman plot for each subregion was plotted in order to visually compare the differences between the models. As shown in Figure 6, the multi-modality model had higher ICC values on each subregion compared to single-modality model, suggesting that multi-modality segmentation was more consistent across longitudinal scans.

**Figure 6.**
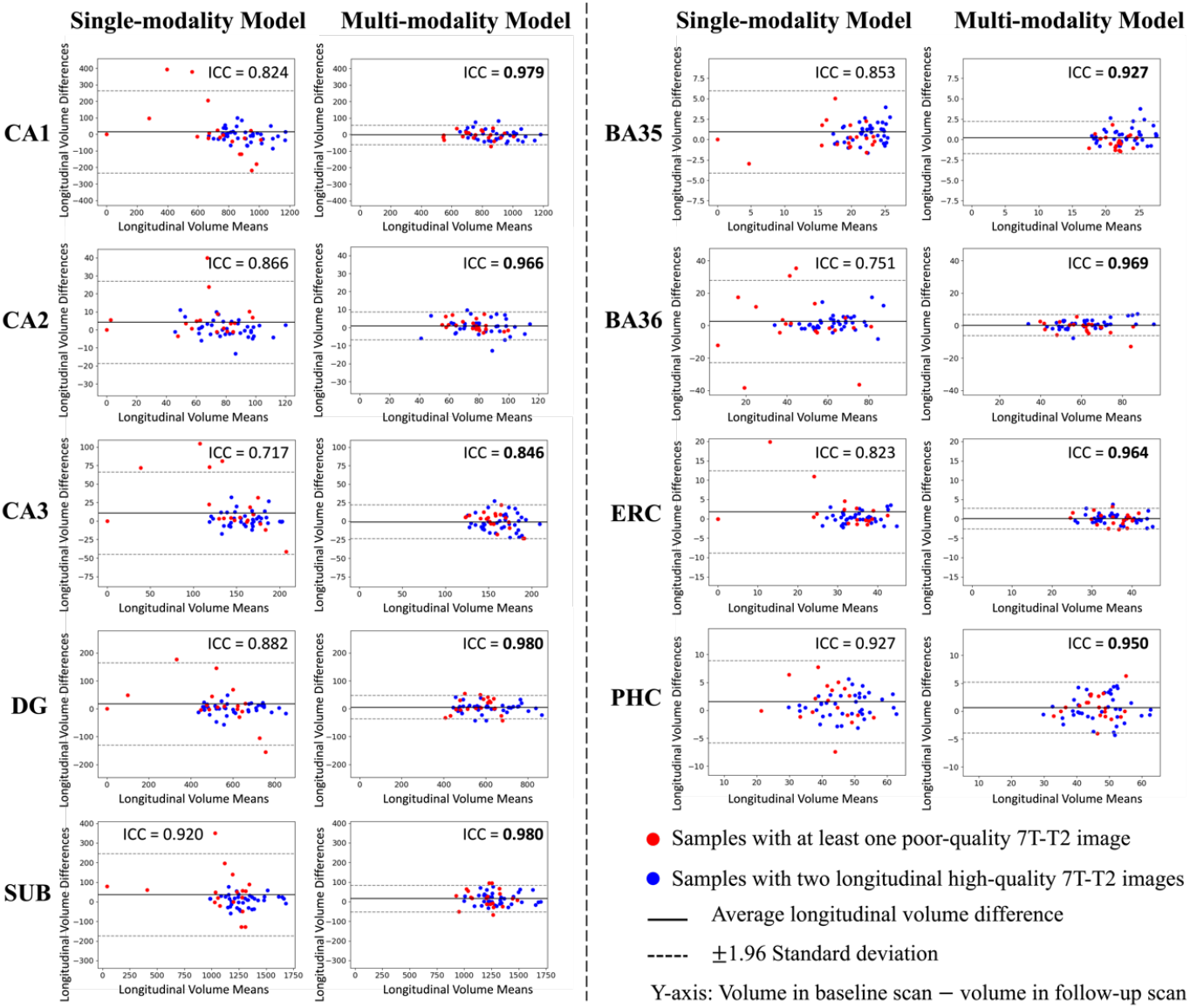
Longitudinal segmentation consistency comparison. For the 29 participants with two longitudinal scans, each scan was segmented using both the multi-modality model and the single-modality model, and the volume of each subregion was calculated from the segmentation. In this figure, the volume consistency between the two scans was represented by Bland-Altman plot and intra-class correlation (ICC) for each model. The multi-modality model had smaller volume mean values and smaller standard deviations of longitudinal volume difference in all subregions. For the single-modality model, the distribution of results from poor-quality data (red dots) was more scattered. . The multi-modality model provided much higher ICC values in all subregions. Higher ICC values are highlighted in bold

As can be seen from the Bland-Altman plot in Figure 6, the segmentation results of multi-modality model in the longitudinal scan pairs had smaller standard deviation, regardless of the quality of the image. The segmentation results of the single-modality model were less consistent across the two scans than those of the multi-modality model. In terms of the distribution of image quality, inconsistency was mainly in the low-quality images.

### 3.5 Discrimination between CU and MCI

The *p*-value, AUC, and Cohen’s D of the fitted GLMs to distinguish A-CU and A+MCI are shown in Table 6. The multi-modality model was associated with a greater difference and effect size between the two diagnostic groups, in absolute terms, compared to single-modality model.

**Table 6.** Amyloid+MCI and Amyloid−CU comparison in the test set. The volumes of hippocampus subfields and median thicknesses of cortical subregions of 65 Amyloid−CU and 18 Amyloid+MCI participants were calculated separately based on the segmentation results of single-modality and multi-modality models. The volumes of subfields were adjusted for the hippocampal tail. p-value, AUC and Cohen’s D of multi-modality model and single-modality model were compared. For each subregion, if the p-value of one of the two models was less than 0.05, the smaller p-value as well as the larger AUC and Cohen’s D were marked bold. (MCI: mild cognitive impairment; CU: cognitively unimpaired; AUC: area under the ROC curve)

| Side | Measure | Subregion | Single-modality Model |  |  | Multi-modality Model |  |  |
| --- | --- | --- | --- | --- | --- | --- | --- | --- |
|  |  |  | *p*-value | AUC | Cohen's D | *p*-value | AUC | Cohen's D |
| Left | Adjusted Volume | CA1 | 0.047 | 0.73 | 0.48 | **0.00040** | **0.75** | **0.86** |
|  |  | CA2 | 0.0098 | 0.69 | 0.78 | **3.0E-05** | **0.78** | **1.25** |
|  |  | CA3 | 0.38 | 0.58 | 0.23 | **0.00029** | **0.76** | **0.93** |
|  |  | DG | 0.057 | 0.71 | 0.49 | **7.4E-05** | **0.78** | **0.99** |
|  |  | SUB | 0.080 | 0.73 | 0.48 | **0.00010** | **0.76** | **0.89** |
|  | Median Thickness | ERC | 0.43 | 0.50 | 0.24 | **0.033** | **0.65** | **0.67** |
|  |  | BA35 | 0.61 | 0.49 | 0.09 | 0.39 | 0.51 | 0.30 |
|  |  | BA36 | 0.28 | 0.41 | 0.26 | 0.71 | 0.46 | 0.01 |
|  |  | PHC | 0.37 | 0.39 | 0.28 | 0.66 | 0.50 | 0.16 |
| Right | Adjusted Volume | CA1 | 0.033 | 0.66 | 0.46 | **0.0012** | **0.72** | **0.66** |
|  |  | CA2 | 0.043 | 0.69 | 0.60 | **0.00062** | **0.77** | **0.94** |
|  |  | CA3 | 0.45 | 0.62 | 0.15 | **0.026** | **0.67** | **0.48** |
|  |  | DG | 0.013 | 0.71 | 0.63 | **0.0010** | **0.73** | **0.78** |
|  |  | SUB | 0.012 | 0.68 | 0.56 | **0.0036** | **0.69** | **0.64** |
|  | Median Thickness | ERC | 0.45 | 0.47 | 0.16 | 0.21 | 0.58 | 0.42 |
|  |  | BA35 | 0.71 | 0.48 | 0.04 | 0.21 | 0.58 | 0.44 |
|  |  | BA36 | 0.54 | 0.45 | 0.10 | 0.91 | 0.52 | 0.14 |
|  |  | PHC | 0.38 | 0.43 | 0.24 | 0.68 | 0.52 | 0.14 |

### 3.6 Generalization performance

The multi-modality segmentation pipeline proposed in this paper was used in a newly released 7T and 3T paired dataset (Chu et al., 2025). This dataset is different from our dataset in various aspects such as 7T scanner, 7T-T1w acquisition protocol, 3T-T2w resolution and age distribution of subjects. Our proposed model successfully finished segmentation on this dataset and the various steps in the pipeline were correctly perpormed. Specific examples can be seen in the Supplementary Material (S9) and complete segmentations are available at our published dataset. This validates the generalization performance of our proposed pipeline on the new dataset.

## 4. Discussion

We developed a multi-modality segmentation model for MTL subregions in structural MRI using 7T-T2w, 7T-T1w (INV1 and INV2), 3T-T2w and 3T-T1w modalities and a modality augmentation scheme to guide the model to learn features from all available modalities.

We found that the multi-modality model trained using the ModAug scheme improved MTL subregion segmentation accuracy slightly over the single-modality 7T-T2 model in cross-validation experiments conducted in high-quality images. However, since the available annotated training data were not representative of the distribution of image quality in the rest of our imaging dataset, we also carried out additional evaluation on downstream tasks, such as comparison of subregion volume (Section 3.3), longitudinal consistency analysis (Section 3.4), and discrimination between the A+MCI and the A-CU groups (Section 3.5). We found that on these downstream tasks, the multimodality model performed similar to the single-modality model on high-quality images, but clearly outperformed it on images of lower quality (Supplementary Material, subsection S4).

Using ModAug encouraged the model to extract information from all available modalities during training. It not only improved the efficiency of the model in utilizing multiple modalities, but also gave the model the ability to cope with modality missing during inference, allowing the use of the model to be not limited to multi-modality data (Supplementary Material, subsection S5, S6 and S7). As a training scheme, it can be combined in future work with other methods for coping with missing modalities, such as image synthesis models (Billot et al., 2023; Pan et al., 2022; Zhou et al., 2022), as explained in subsection S6 of the Supplementary Material.

Furthermore, with the ability of the model to cope with missing modality, we explored the contribution of different modalities in the multi-modality model and found that the introduction of 3T images helped to improve the segmentation accuracy (as shown in subsection S6 of the Supplementary Material), which reinforced the importance of multi-modality data across the MRI fields for designing robust segmentation algorithms.

In the following section, we discuss the differences between our proposed model and other existing works in terms of methodology, evaluation metrics, and results.

### 4.1 Methodological differences from the works of other research groups

Table 7 is the summary of existing automatic segmentation methods for hippocampal subfields or MTL subregions.

**Table 7.**
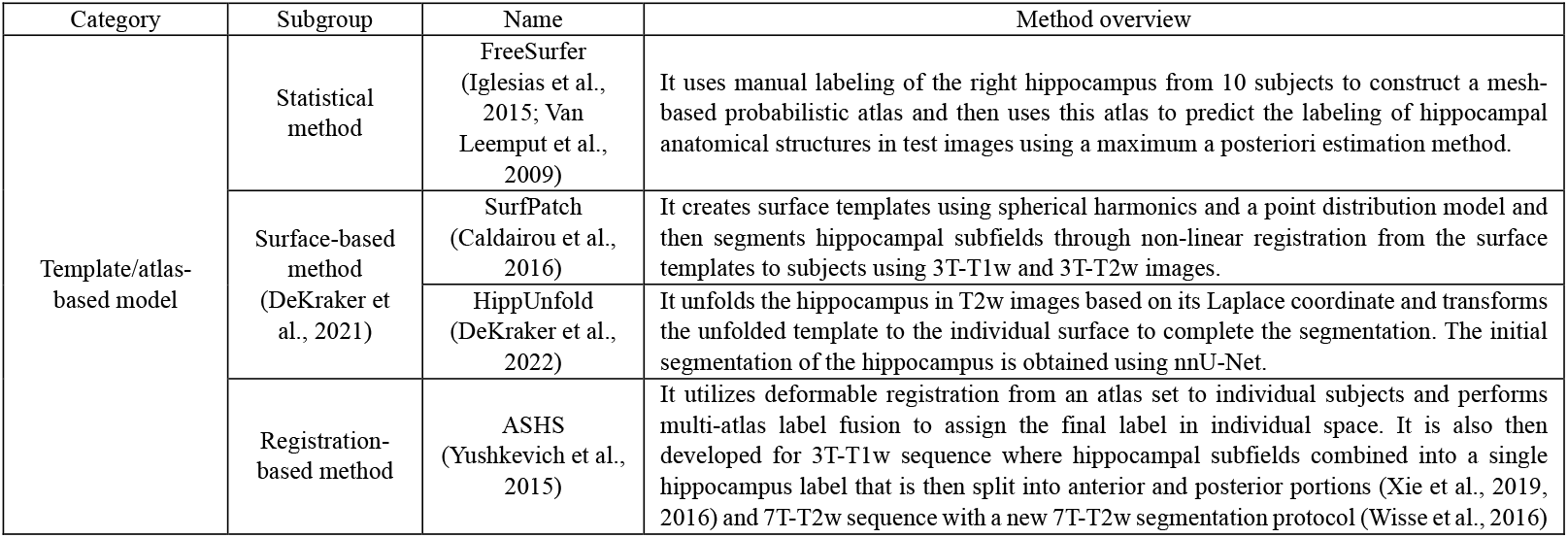
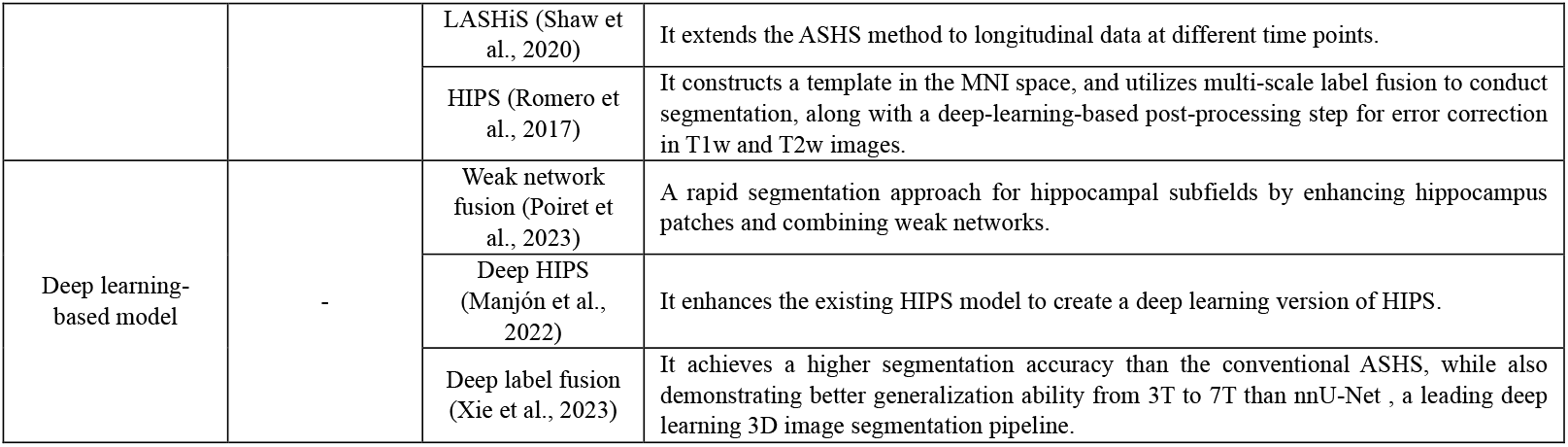
The overview of existing automatic segmentation for hippocampal subfields and MTL subregions.

Building upon the foundation of the ASHS framework, our work presents differences over both the traditional ASHS and other segmentation methods listed above.

When compared with atlas/template-based methods, which explicitly utilize atlas information, our proposed deep learning-based model implicitly encodes this information within the neural network’s weights. This is achieved through the optimization process during training. According to findings by (Xie et al., 2023), learning-based MTL segmentation approaches can yield higher accuracy within identical data domains but may exhibit reduced generalizability across image domain.

When compared with other deep learning-based methods, our proposed approach necessitated additional registration steps, particularly the inter-field alignment between 3T and 7T images. This ensured that the superior image qualities of different modalities are effectively integrated into the primary modality, but might introduce registration errors .

Regarding the original intent in designing models, our method has one important difference compared to other methods: we did not intend to propose a state-of-the-art subregion segmentation method that can surpass all existing models. Instead, our objective was to verify the feasibility of utilizing multi-modality data to compensate the deficiency in segmentation precision of single-modality models in the context of low-quality images. The majority of existing methods are single-modality or single-field models and the idea of integrating multi-modality data has the potential to enhance the performance of these existing models.

Among the existing methods, HippUnfold is one of the most recently proposed and has been evaluated on 7T-T2w images (DeKraker et al., 2022). Therefore, we repeated our longitudinal consistency experiments using HippUnfold to demonstrate that single-modality models are prone to the common issue of reduced accuracy in low-quality images. The longitudinal 7T-T2w MR images from the 29 subjects used in section 3.4 were segmented using trained HippUnfold by the authors, and then the longitudinal consistency of the subfield volume was evaluated. The Bland-Altman plots and ICC values are presented in Figure 7.

**Figure 7.**
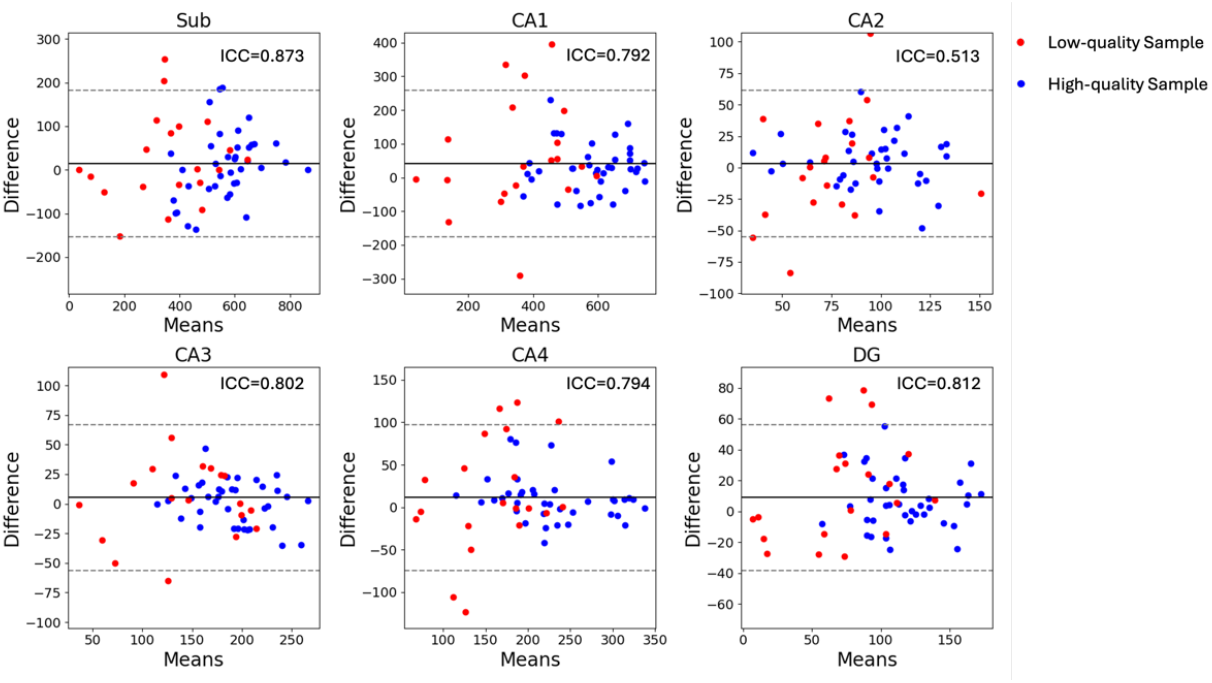
The Bland-Altman plots for the longitudinal volumes of hippocampal subfields segmented by HippUnfold. Dashed line: 1.96 standard deviation.

From Figure 7, it is evident that the HippUnfold exhibited lower consistency (higher variance and a more dispersed distribution) on low-quality compared to high-quality images. This was consistent with our findings for the single-modality nnU-Net model in section 3.4. This is not surprising, since HippUnfold itself uses a single-modality nnU-Net to generate an initial segmentation and the segmentation is applied in surface fitting steps. The results of this study suggest that the performance of HippUnfold on 7T-T2 could also be improved by incorporating other 7T and 3T modalities with the ModAug strategy into this initial segmentation step.

### 4.2 Different segmentation evaluation metric

When evaluating the accuracy of a segmentation model, it is common to report average metrics such as Dice coefficient in cross-validation experiments conducted on all available annotated data. However, there may be an inherent bias in image quality between images that are annotated for training and images on which the trained model is ultimately used. This was indeed the case in our dataset, where the proportion of low-quality images was much higher for the non-annotated test images (32.85% in test set vs 6% in training set). If we only considered this type of evaluation, we would have concluded that there is no benefit from using multiple modalities in MTL subregion segmentation. The Dice coefficient computed in cross-validation experiments did not reflect the superior performance of multi-modality model because the data in the training set were selected from high quality images, and as demonstrated in Table 4 and Figure 5. The single-modality model also obtained good segmentation results from high-quality images.

However, our indirect evaluation on the non-annotated test set clearly points to greater robustness of the multi-modality model. In order to compare the models’ performance in images with different qualities, we evaluated them in a test dataset that included low quality images. However, manual annotation of these data was not practical, or outright impossible, precisely because the image quality of the modality used for manual segmentation was low. Hence, quantitative analyses, including measuring Dice coefficient, could not be performed. Instead, we measured the robustness of the segmentations generated by the two models in images with different qualities through a series of indirect tasks, where two aspects were verified: stability and accuracy. Specifically, by comparing the segmentation results of the two models and by comparing the segmentation results of the same model at different longitudinal time points, we verified that the multi-modality model likely had better segmentation stability, particularly in low-quality images. By demonstating greater separation between disease and control groups (higher AUCs) on a discrimination task, we also got strong evidence in the higher segmentation accuracy of the multimodality model.

### 4.3 Differences in experimental results from prior work

When distinguishing between A+MCI and A-CU, difference in median thickness for BA35 was not significant and in ERC thickness was only significantly smaller on the left. This was inconsistent with our experience and with existing research (Duara et al., 2008; Hata et al., 2019; Ogawa et al., 2019; Wolk et al., 2017; Xie et al., 2020; Yushkevich et al., 2015), especially given that the volumes of hippocampus subfields were significantly smaller in A+MCI group. Since BA35 approximates the transentorhinal region, the site of earliest cortical tau tangle pathology and neurodegeneration in AD, we expect this region to show differences as pronounced as the hippocampus and hippocampal subfields. Earlier MTL morphometry studies using ASHS also showed results consistent with these expectations (Wolk et al., 2017; Yushkevich et al., 2015).

We identified four possible reasons for this discrepancy. First, the definition of BA35 boundaries in the current protocol may not completely align with prior versions of procotol (Yushkevich et al., 2015) used in prior studies. Second, the segmentation by nnU-Net may be less accurate (oversegmentation or undersegmentation) than ASHS. Third, the small sample size (18 A+MCI) caused the lack of significant effect. Fourth, the image resolution used in this study differs from that of previous studies, which may affect the accuracy of the thickness calculation.

To find the underlying causes of the observed discrepancies, we undertook a series of supplementary experiments. In addition to employing the 7T-T2w atlas developed using the ABC protocol, we leveraged all available atlases within our institution to train segmentation models. Subsequently, we evaluated their performance on discrimination tasks between A-CU and A+MCI cohorts, thereby facilitating a comparative analysis across various imaging resolutions, protocols, and primary modalities. The additional atlas sets utilized for this purpose comprised: 1) a 3T-T1w atlas constructed in accordance with the Penn Memory Center (PMC) protocol (Xie et al., 2019), wherein 3T-T1w images underwent preprocessing via non-local means (Manjón et al., 2010) to generate images super resolution; 2) a 3T-T2w atlas also based on the PMC protocol (Yushkevich et al., 2015); and 3) a separate 3T-T2w atlas derived from the ABC protocol.

Three additional nnU-Net models were trained using three different atlas sets, and an existing ASHS model was trained using the 3T-T1w PMC atlas. These models were then tested on the same test set used in this study. The p-values of the GLM in discriminating between A-CU and A+MCI are presented in Table 8, along with the protocol names and primary test resolution.

**Table 8.**
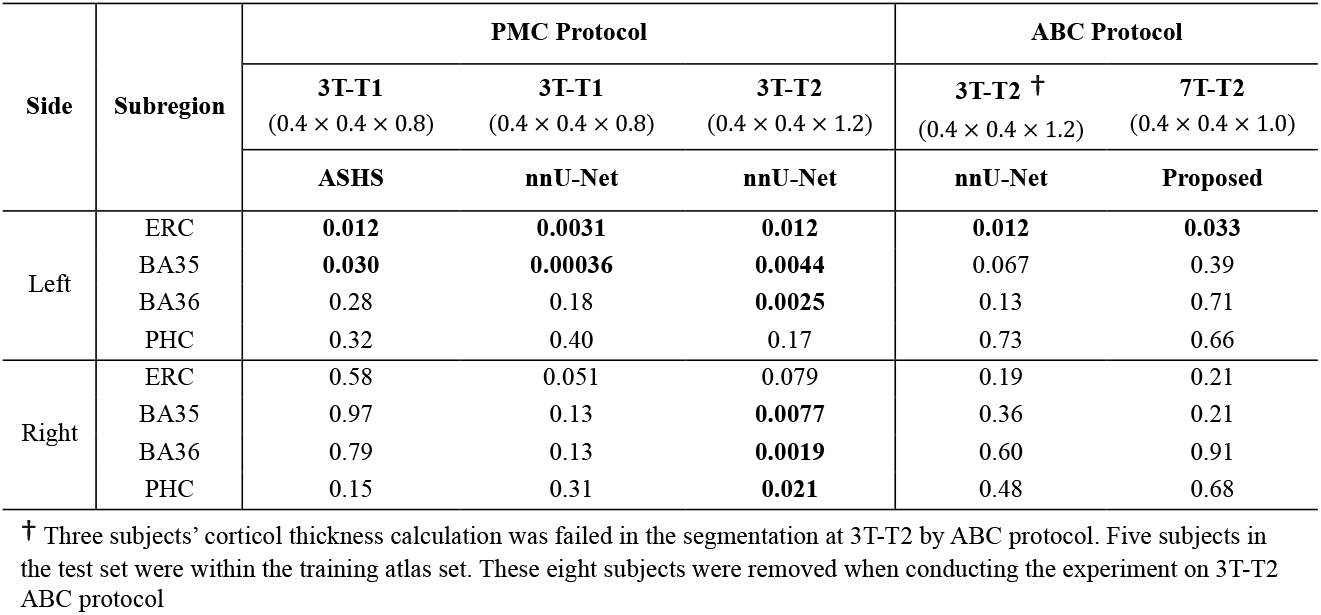
The p-value comparisons of subregion thickness in A-CU and A+MCI discrimination among different primary modalities, resolutions and protocols.

| Side | Subregion | PMC Protocol |  |  | ABC Protocol |  |
| --- | --- | --- | --- | --- | --- | --- |
|  |  | 3T-T1<br>(0.4 × 0.4 × 0.8) | 3T-T1<br>(0.4 × 0.4 × 0.8) | 3T-T2<br>(0.4 × 0.4 × 1.2) | 3T-T2 †<br>(0.4 × 0.4 × 1.2) | 7T-T2<br>(0.4 × 0.4 × 1.0) |
|  |  | ASHS | nnU-Net | nnU-Net | nnU-Net | Proposed |
| Left | ERC | **0.012** | **0.0031** | **0.012** | **0.012** | **0.033** |
|  | BA35 | **0.030** | **0.00036** | **0.0044** | 0.067 | 0.39 |
|  | BA36 | 0.28 | 0.18 | **0.0025** | 0.13 | 0.71 |
|  | PHC | 0.32 | 0.40 | 0.17 | 0.73 | 0.66 |
| Right | ERC | 0.58 | 0.051 | 0.079 | 0.19 | 0.21 |
|  | BA35 | 0.97 | 0.13 | **0.0077** | 0.36 | 0.21 |
|  | BA36 | 0.79 | 0.13 | **0.0019** | 0.60 | 0.91 |
|  | PHC | 0.15 | 0.31 | **0.021** | 0.48 | 0.68 |

† Three subjects' cortical thickness calculation was failed in the segmentation at 3T-T2 by ABC protocol. Five subjects in the test set were within the training atlas set. These eight subjects were removed when conducting the experiment on 3T-T2 ABC protocol

Analysis of Table 8 yields three observations:

1. When comparing the segmentation performance of ASHS with nnU-Net on 3T-T1w images, nnU-Net demonstrated higher significance. This suggests that nnU-Net had robust subregion segmentation capabilities and was unlikely to be a contributing factor to the noted discrepancy.
2. Given that all experiments were conducted using the same dataset and that certain experiments revealed significant findings pertaining to BA35 thickness, it can be inferred that the dataset itself did not account for the observed discrepancy.
3. Upon examining both imaging protocols and resolutions, it appeared that protocol might play a more substantial role than image resolution in driving the discrepancy. This was evidenced by the fact that all left BA35 thickness measurements derived from PMC protocol consistently exhibited statistical significance (p<0.05) when discriminating between A-CU and A+MCI, whereas those obtained from ABC protocol did not achieve such significance. Additionally, the 3T-T2 PMC atlas achieved significant results in most left and right MTL subcortical structures, whereas 3T-T2 ABC atlas did not, again suggesting that the protocol, rather than resolution, is the source of the discrepancy.

According to the original publications for PMC protocol (Yushkevich et al., 2015) and ABC protocol (Berron et al., 2017), these protocols employ slightly different rules to label MTL subregions and these differences might explain the lack of group effects in BA35 and right ERC, even though they were developed in collaboration with the same neuronatomist (Ding and Van Hoesen, 2010).

To demonstrate these differences, two participants’ segmentation of BA35 in 7T-T2w by multi-modality model trained by ABC protocol and 3T-T1w by nnU-Net trained by PMC protocol are shown in Figure 8. For the first participant in Figure 8, BA35 was located on the medial side (i.e., left) of the collateral sulcus in both 3T-T1w and 7T-T2w. For the second individual, the BA35 in 3T-T1w was mainly located on the medial side of the collateral sulcus and BA35 in 7T-T2w was on medial side in regions when the the sulcus was deep (the first slice in coronal view) and on both sides when more shallow (the second and third slices in coronal view).

**Figure 8.**
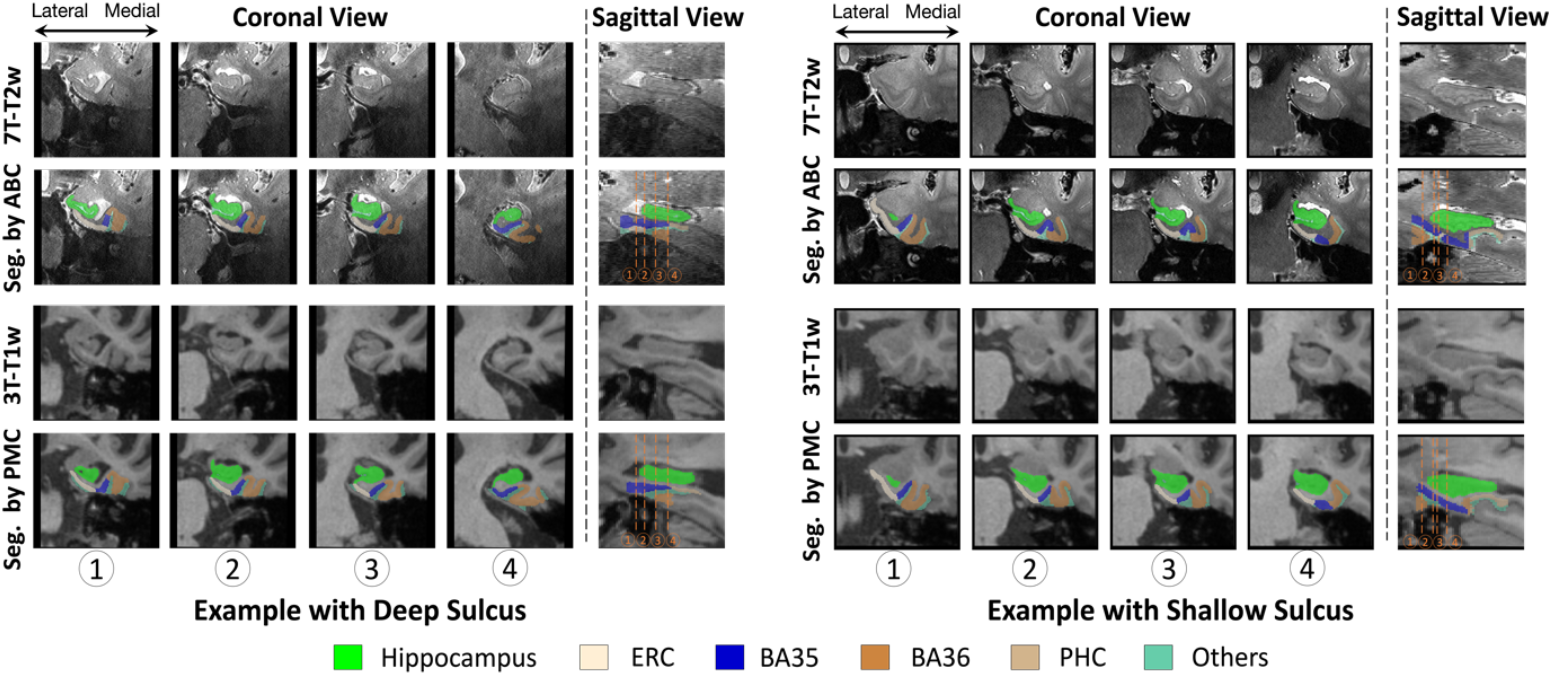
Comparison of BA35 (dark blue) segmentation by nnU-Net trained in PMC protocol (3T-T1w) and multi-modality model trained in ABC protocol (7T-T2w). The first example had deep sulcus and segmentations for BA35 by both methods were at the medial side of collateral sulcus. The second example had shallow sulcus in slice 2 and 3 in coronal view and segmentations of BA35 by multi-modality model were on both sides of collateral sulcus. (The 3T-T1w images and segmentations have been registered to 7T-T2w here, making it possible to compare them with segmentations in 7T-T2w.)

However, we have only verified that the difference in protocols was a possible reason for the difference in significance in the experiments, but have not found a specific way of how exactly the ABC protocol affects the thickness of the subregion. Further experiments are necessary in the future.

### 4.4 Cost-effectiveness

One of the main benefits from our multi-modality model is that it increases the effectiveness of research that uses multi-scanner/multi-modality MRI and relies on MTL subregional segmentations for quantitative analysis. For example, at our Alzheimer’s disease research center, 3T and 7T MRI is acquired in the same participants due to complementary strengths of these scanners, with the 3T scanner configured to provide superior diffusion-weighted imaging, and 7T allowing higher resolution for structural imaging and better imaging of myelin and iron-related pathology (Brown et al., 2024).

As shown in Table 5, by only using a single-modality model, a significant proportion of MTL segmentations 30.81% (or 10.47%, if using a less stringent quality threshold) would be rejected due to low quality. With the multi-modality approach, only 12.79% (or 0.29%) would be rejected, representing a 58.49% (or 97.22%) reduction in the number of rejected segmentations. This represents a significant improvement in our ability to utilize both MTL subregional morphometry and downstream measures extracted from other 7T modalities, and a significantly more effective use the volunteer participants time and effort.

### 4.5 Limitations and future work

The goal of modality augmentation is to force the model during training to allocate attention to all available channels and not focus too much on the primary modality, which in the high-quality training set is likely to be most useful for minimizing the network loss but is more likely to have low-quality in datasets encountered at inference. Filling certain input channels with Gaussian noise is a specific implementation of modality augmentation, and other implementations are possible. For example, filling channels with zero instead of Gaussian noise achieved similar performance in preliminary experiments. More advanced augmentation strategies that synthesize realistic low-quality images during training may achieve better performance; however, since there are many possible synthesis strategies (diffusion, adversarial models, physics-based MRI artifact simulation), we felt that this approach is best explored in future work.

Another limitation is that the proposed multi-modality model required one modality to be selected as “primary”, in our case this was 7T-T2w. This modality is used as a target for registration and all other modalities are resampled in its space, causing loss of advantage of other modalities’ resolution, such as the isotropic resolution and finer plane thickness in 3T-T1w and 7T-T1w. This may be one of the factors affecting thickness calculations, which in turn affects the accuracy of downstream tasks, such as distinguishing between A+ MCI and A-CU. In the future, it may be possible to develop a model that segments each modality in its own space while incorporating inter-information from others. This would optimize data availability and maintain each modality’s resolution advantages.

In our future work, in addition to the two ideas mentioned above, we will explore various combinations of modalities for model training and examine how the image quality of different modalities affects segmentation results during inference as a supplement to this paper.

## 5. Conclusion

This study developed a segmentation model for MTL subregions using 7T-T2w, 7T-T1w, 3T-T2w and 3T-T1w MR images. Incorporating these modalities during training with the help of together with modality augmentation led to a model that is more resilient to low image quality, resulting in more accurate segmentation. When the primary modality’s image quality was low, proposed multi-modality model still could generate stable segmentaitons by extracting useful information from other modalities.

While the current study focused on a very unique dataset with four structural modalities collected at two MRI field strengths, the challenges and the insights gained in the evaluation regarding the utility of multi-modality models in the context of poor/variable image quality and the danger of only using cross-validation performance to select the best segmentation model when manually annotated data exhibit selection bias relative to the real-world data distribution are likely relevant to other image segmentation contexts with multiple modalities available, such as DTI, SWI.

## Author Contributions

**Yue Li**: Conceptualization, Formal analysis, Investigation, Methodology, Validation, Visualization, Writing – original draft. **Long Xie**: Data curation, Methodology, Writing – review and edition. **Pulkit Khandelwal**: Methodology, Writing – review and edition. **Laura E. M. Wisse**: Data curation, Resources, Writing – review and edition. **Christopher A. Brown**: Data curation, Resources. **Karthik Prabhakaran**: Resources. **M. Dylan Tisdall**: Resources, Writing – review and edition. **Dawn Mechanic-Hamilton**: Resources. **John A. Detre**: Data curation, Project administration, Resources, Writing - review and editing. **Sandhitsu R. Das**: Conceptualization, Data curation, Formal analysis, Investigation, Methodology, Project administration, Resources, Supervision, Writing - review and editing. **David A. Wolk**: Conceptualization, Data curation, Formal analysis, Investigation, Methodology, Project administration, Resources, Supervision, Writing - review and editing. **Paul A. Yushkevich**: Conceptualization, Data curation, Formal analysis, Investigation, Methodology, Project administration, Resources, Supervision, Funding acquisition, Writing - review and editing.

## Consent for publication

All authors have reviewed the contents of the manuscript being submitted, approved of its contents and validated the accuracy of the data and consented to publication.

