## supplemental materials for "Automatic segmentation of medial temporal lobe subregions in multi-scanner, multi-modality magnetic resonance imaging of variable quality"

### Supplementary Material

#### S1. Inclusion and exclusion criteria

The inclusion and exclusion criteria used in this study are as follows.

**Inclusion criteria**:

- Participants with 7T and 3T MRI scanned within two years.

**Exclusion criteria**:

- Participants who did not have a diagnosis of CU or MCI at the most recent diagnosis to the date of the 7T MRI scan.
- Participants whose diagnosis results closest to the 3T MRI scan are inconsistent with the diagnosis results closest to the 7T MRI scan.
- Participants with at least one missing modality in 7T-T1w, 7T-T2w, 3T-T1w and 3T-T2w.
- Participants who experienced registration failure during registration (described in section 2.2) from other modalities to 7T-T2w.

To collect longitudinal scans for test participants, one inclusion criteria and one exclusion criteria were added to the criteria above. The **additional inclusion criteria** is "The participants who had more than one 7T/3T pairs" and the **additional exclusion criteria** is "The participants whose diagnoses in two longitudinal 7T scans were not consistent".

Note 1: There was one exception in the training set that did not meet the inclusion criteria. The time interval between the 7T and 3T scans in this case was 33 months, and the most recent diagnosis on the 7T scan (the earlier scan) was MCI, and the most recent diagnosis on the 3T scan (the later scan) was CU.

Note 2: As explained in the Discussion, the proposed model has the ability to process the situation of missing modality as long as 7T-T2w image is available. The reason why we excluded the subject with missing modality was that we need to compare the difference between the model with and without full modalities as input.

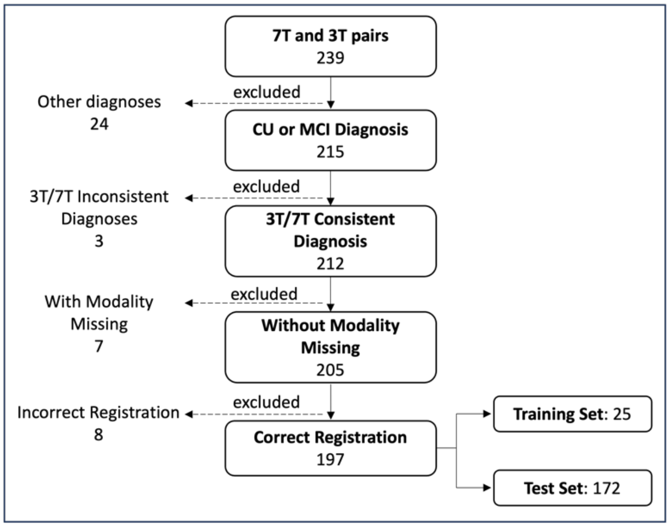


*Figure S1 Participant selection according to inclusion and exclusion criteria.*

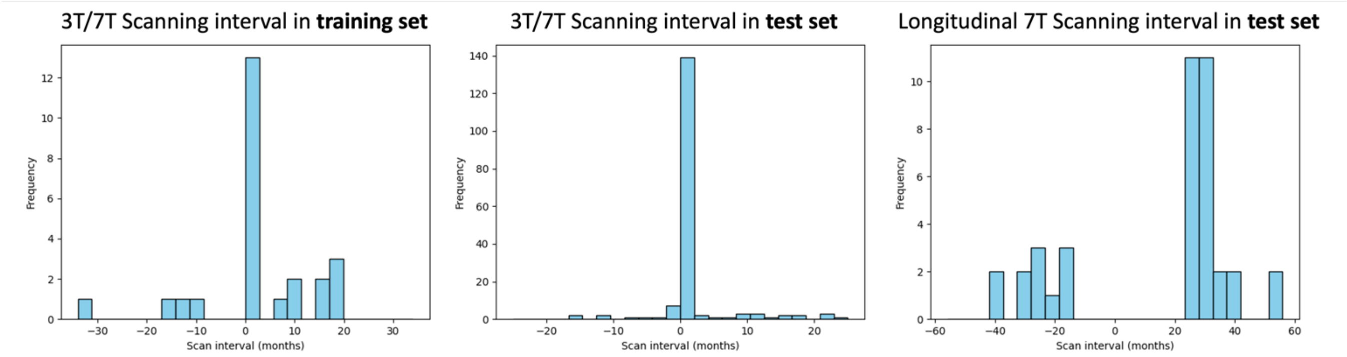


*Figure S2 distributions of Scanning interval between 3T and 7T MRI scans in both training and test set and longitudinal 7T scans in test set*

### S2. Example of visual registration checking

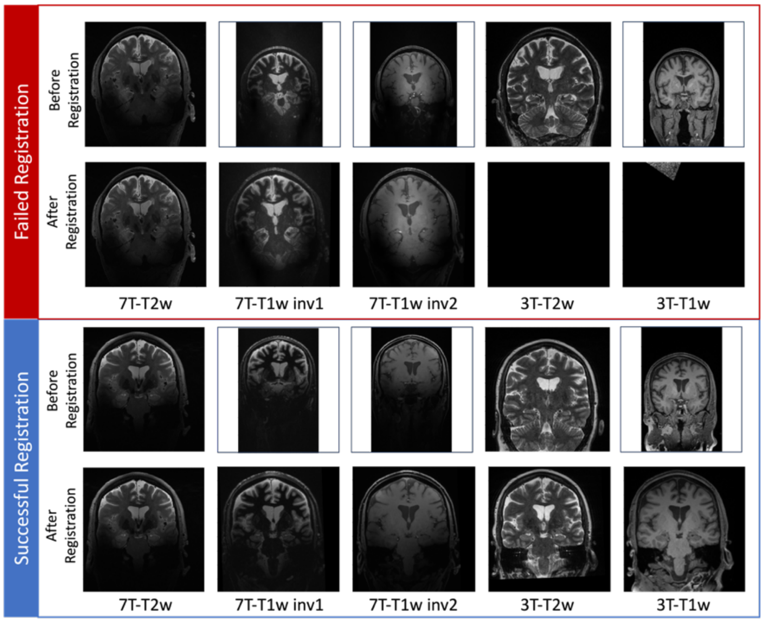


*Figure S3 Examples of failed and successful whole-brain registration. In the first example, the registrations from 3T-T1w and 3T-T2w to 7T-T2w were failed. This situation was excluded by visual checking.*

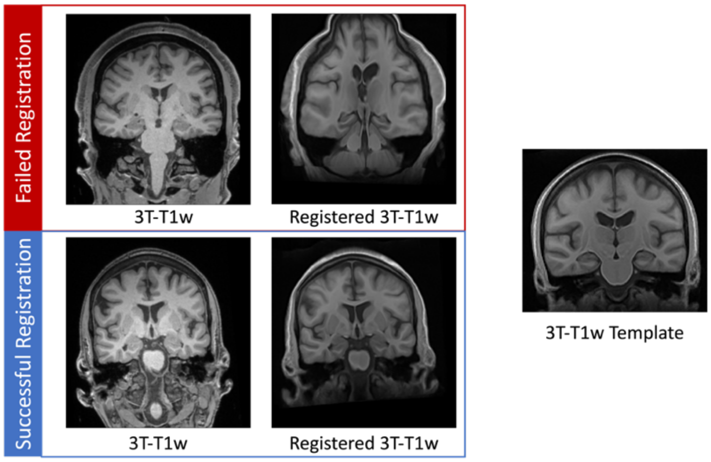

*Figure S4 Examples of failed and successful 3T-T1w template deformable registration. In the first example, the registrations from 3T-T1w template to 3T-T1w was failed. This situation was excluded by visual checking.*

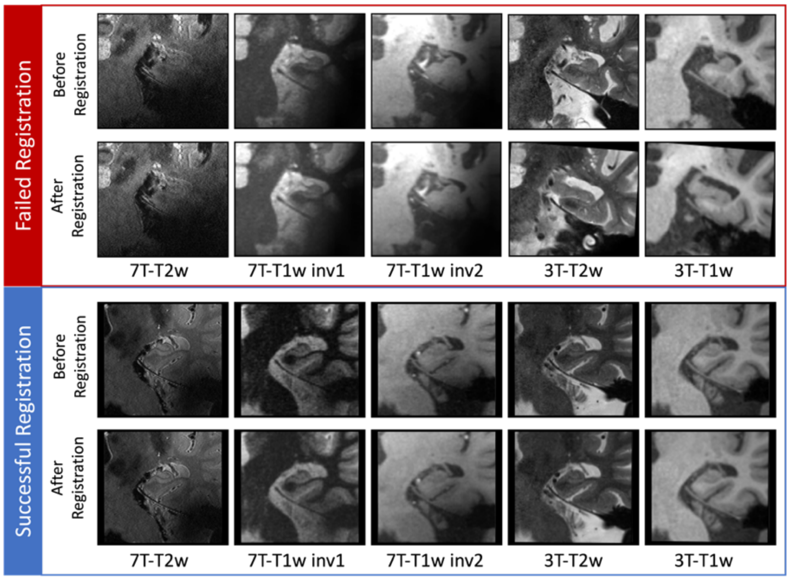


*Figure S5 Examples of failed and successful local registration. In the first example, the registrations from 3T patches to 7T-T2w patch was failed. This situation was excluded by visual checking.*

### S3. Schematic diagram of data augmentation

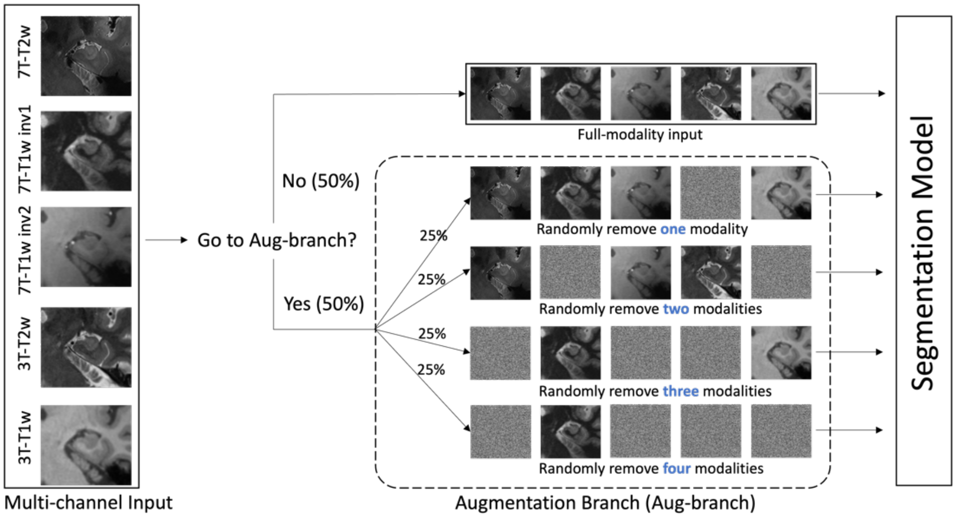


*Figure S6 The pipeline of modality augmentation. When five channels were fed into the network, they would be processed by augmentation branch with probability of 50%. Othewise, the full modalities were used in the segmentation. In augmentation branch, there was equal probability that one, two, three and four modalities were replaced by noise randomly.*

### S4. Results of removing outliers

Based on our findings from section 3.3 and section 3.4, we noticed that the segmentation performance of single-modality and multi-modality models differs mainly due to poor-quality images. We anticipate that if we eliminate these poor-quality samples, both models should demonstrate similar performance in both segmentation and downstream tasks.

To achieve this, we employed various thresholds (0, 1, 2, 3, and 4) to identify outliers. In each iteration, images with a quality rating below the corresponding threshold were excluded, and we assessed the longitudinal consistency and the segmentation's ability to differentiate between A-CU and A+MCI at each threshold. The results are illustrated in Figure S7, showing the ICC for longitudinal consistency, and Figure S8, presenting the AUC in GLM for discriminating between A-CU and A+MCI.

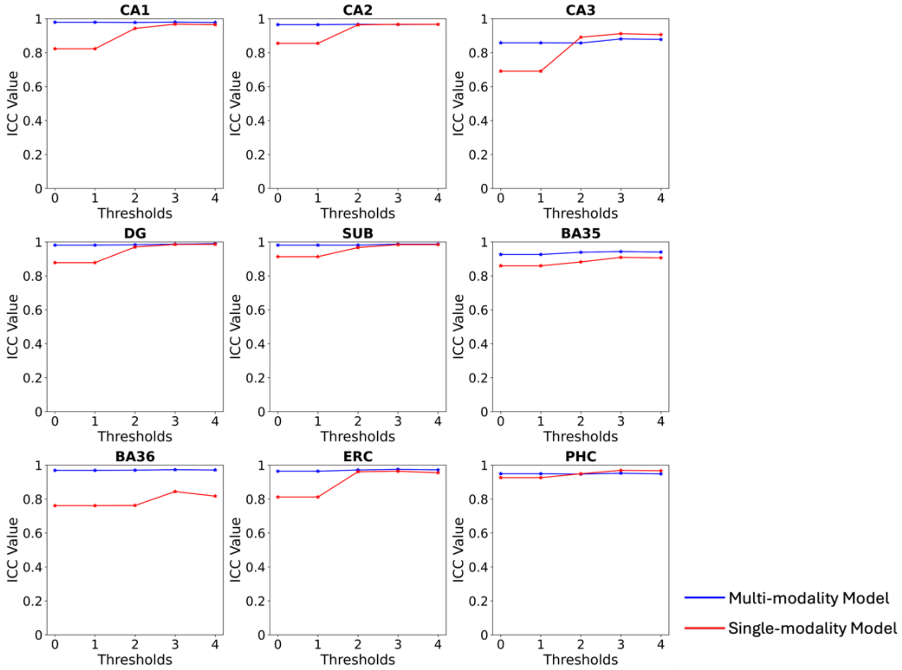


*Figure S7 ICC values comparison across subregion volumes under each threshold to remove outliers in single-modality and multi-modality models.*

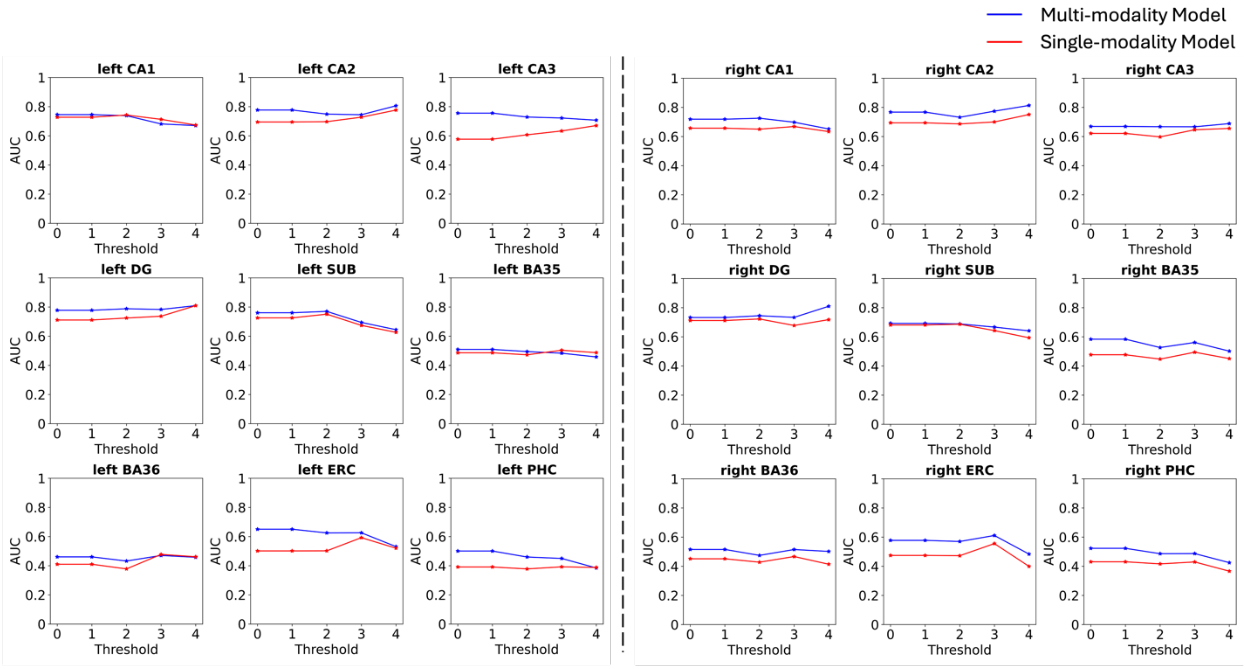


*Figure S8 The AUC comparison of single- and multi-modality models in image samples at each threshold.*

In Figure S7, the ICC values of the single-modality model increase as the threshold increases, while the ICC values of the multi-modality model remain consistently high. The ICC values of the two models become very similar when all poor-quality images are removed, except for BA36.

In our discrimination experiment, we separated the left and right sides. In Figure S8, we noticed that as more outliers are removed on the left side, the performance of the two models becomes increasingly similar. This suggests that poor image quality may be contributing to the differing performances of the two models in the discrimination task. However, this trend is not observed on the right side. Previous studies have suggested that the right side is less significant than the left in distinguishing possible AD subjects.

One possible explanation for this phenomenon could be that as we systematically removed outliers from our analysis, we observed a growing imbalance in the ratio between A-CU and A+MCI. At a threshold of 4, for example, the counts for A-CU and A+MCI were 39 and 7. However, when we adjusted the threshold to 0, these figures shifted to 18 for A-CU and 65 for A+MCI, respectively. This trend suggests that the relatively low number of A+MCI cases might make the AUC value unstable and more vulnerable to minor fluctuations, although other factors may also contribute to this.

### S5. The role of modality augmentation

In cross-validation, the multi-modality model with ModAug did not show better segmentation Dice on every subregion than that without ModAug, although they were close enough to. We expected that ModAug would have greater advantage in the test set. The subregion volume comparison of the multi-modality model trained with and without ModAug is shown in Figure S9.

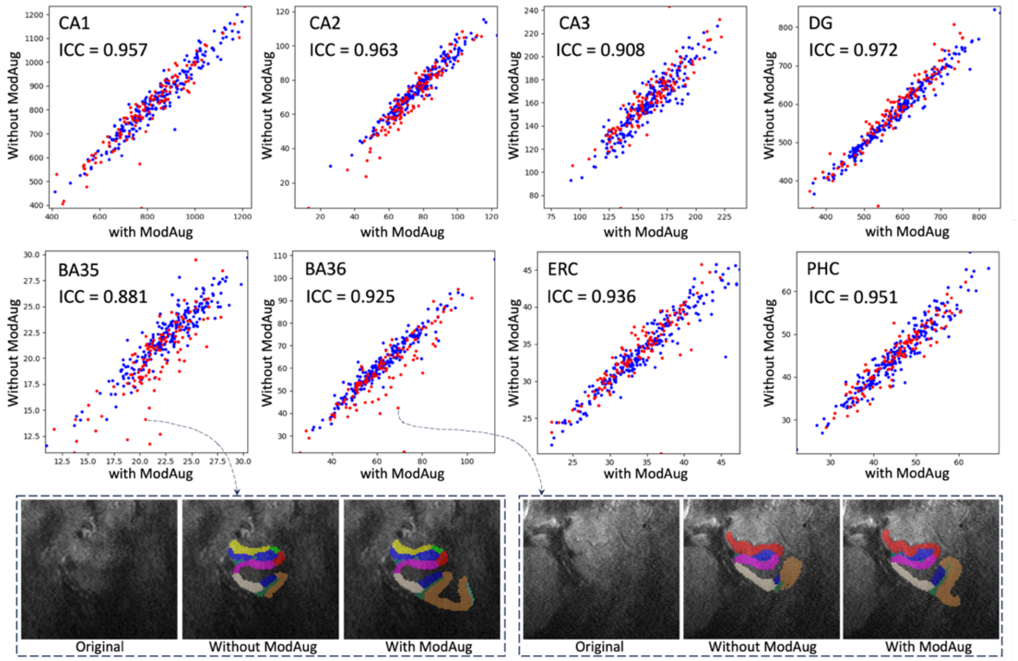


*Figure S9 Subregion volumes comparison between multi-modality model with and without ModAug. (**Red dots**: low-quality images; **blue dots**: high-quality images)*

In Figure S9, except for BA35 and BA36, high-quality dots and low-quality dots have similar distribution in the plots. In BA35 and BA36, there were more poor-quality images as outliers. Two outliers were picked and shown at the bottom of Figure S9. In these two examples, the model trained without ModAug might have undersegmentation in BA35 and BA36. The model trained with ModAug output the results with more reasonable shape of MTL. This is in line with a common issue of signal drop out in more lateral regions in the primary modality 7T-T2w which mostly affects segmentation accuracy of BA35 and BA36. Relying more on the other modalities, which are less prone to signal drop out in these areas, should indeed provide a more complete segmentation of BA35 and BA36.

### S6. Importance of each modality

ModAug appeared to allow extraction of useful information from alternative modalities when image quality was poor in the primary one. It allowed the model to be robust to missing modalities, but we did not know how much information provided by each modality.

In order to further verifiy the importance of each modality, we ran the inference of the multi-modality model with certain modalities discarded. Specificly, 3T, 7T-T1w and both 3T and 7T-T1w were replaced by noise images in the inference, respectively, and we observed the changes of segmentations after removing these modalities.

The subregion volume consistency between these results and the model with complete input (no modality missing) was calculated by ICC. As shown in Table S1, when both 3T and 7T-T1w were missing, there was the most significant loss of information resulting in lower ICC values compared with input in which only one of these modalities was missing. The input missing 7T-T1w had higher ICC values than the input missing 3T, indicating that the model was more dependent on the 3T modality than the 7T-T1w. This conclusion can be also reflected in Figure S10, which shows two poor-quality examples segmented by the multi-modality model on input with different modalities missing. The results on input only missing 7T-T1w were closest to the complete input.

*Table S1 ICC values of subregion volume comparison between multi-modality model with certain modalities missing and model without modality missing*

| Model | Missing modality | CA1 | CA2 | CA3 | DG | SUB | BA35 | BA36 | ERC | PHC |
|---|---|---|---|---|---|---|---|---|---|---|
| **Multi-modality Model with ModAug** | **3T** | 0.995 | 0.989 | 0.981 | 0.972 | 0.992 | 0.971 | 0.98 | 0.988 | 0.986 |
| | **7T-T1w** | 0.998 | 0.994 | 0.99 | 0.997 | 0.998 | 0.986 | 0.998 | 0.993 | 0.992 |
| | **3T and 7T-T1w** | 0.871 | 0.811 | 0.793 | 0.869 | 0.854 | 0.758 | 0.767 | 0.851 | 0.804 |
| **Single-modality Model** | **3T and 7T-T1w*** | 0.795 | 0.747 | 0.662 | 0.783 | 0.75 | 0.696 | 0.675 | 0.595 | 0.786 |

* The single-modality model only used 7T-T2w as input. Therefore, there was no 3T or 7T-T1w fed into the model. We also call this situation as "missing 3T and 7T-T1w" here.

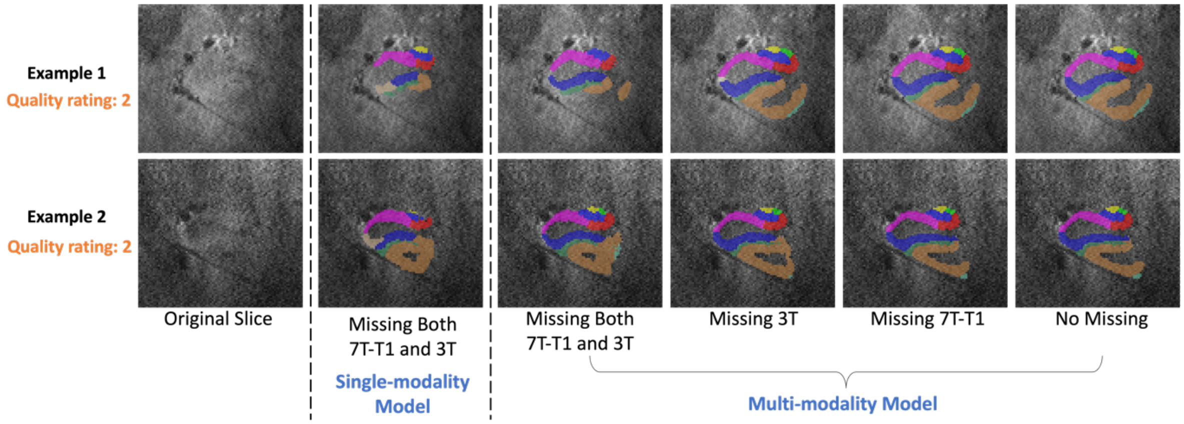


*Figure S10 Two examples of poor-quality images segmented by multi-modality models with input missing 7T-T1w and 3T, missing 3T, missing 7T-T1w and input without modality missing. The segmentations by single-modality for these two examples are also compared here.*

Besides different situations of missing modality, the segmentation by single-modality model was also compared with other scenarios. The ICC values in Table S1 show that single-modality model had lowest consistency with multi-modality model with full modalities as input. The examples in Figure S10 also reflect the low consistency between the single-modality model and multi-modality model. Although we input the same amount of information into single-modality model and multi-modality model with both 3T and 7T-T1w input missing, the single-modality model had more undersegmentation. The reason might be in the training process. For multi-modality model, the information of different modalities from training set was stored in the network parameters of nnU-Net, and when encountering missing modality, these embedded information can help the model make relatively better predictions.

Although ModAug facilitates training multi-modality model to handle inputs when certain modalities are absent, its initial conception was to compel the extraction of information from all accessible modalities, rather than relying solely on the primary one. Other methodologies, such as image synthesis models (Billot et al., 2023; Pan et al., 2022; Zhou et al., 2022), also offer solutions for addressing missing modalities in multi-modality segmentation tasks. A future research involves integrating synthesis model, which has the potential to enhance image quality and replenish missing modalities, with ModAug, thereby advancing our approach to challenges within multi-modality application contexts.

### S7. Feasibility of running inference in non-multimodal data

The proposed multi-modality model involved both 7T and 3T MRI. This raises a question if this model can still be used when these modalities are not all available. After all, there are not many subjects who have all these all modalities. If a model is only useful for multiple modalities, it will limit its applicability.

From the comparisons in section S6, we found this multi-modality model was able to segment MTL subregions using only 7T-T2w modality. Table S1 and Figure S10 indicate that even if only uses single modality, 7T-T2w, the multi-modality model has a better performance than single-modality model, with the full-modality model as the silver standard. This demonstrates that the multi-modality model is compatible with the single-modality model and therefore does not limit the application of it to other datasets without full modalities.

The examples in Figure S10 have low image quality. Figure S11 shows examples of segmentation with good image quality even when certain modalities are missing. The segmentation results are very consistent with or without the missing modality.

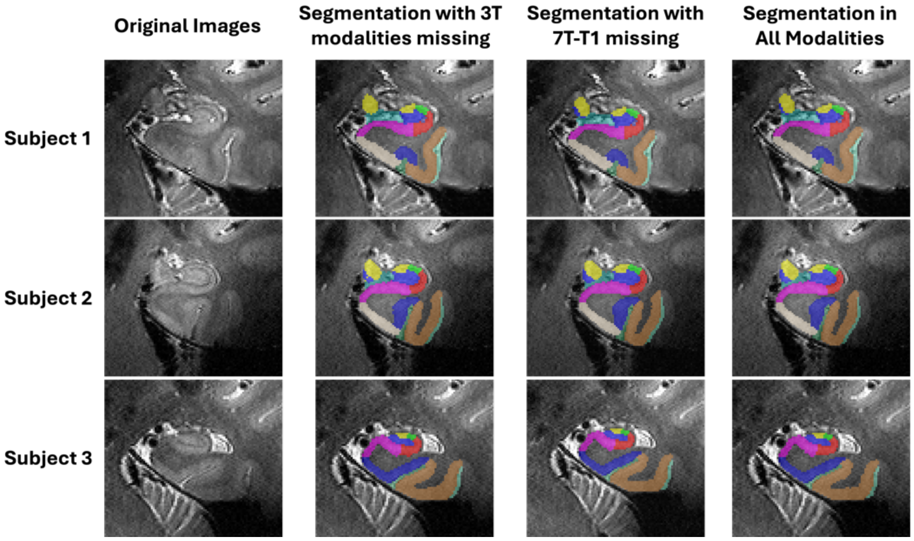


*Figure S11 Segmentation examples of proposed method with 7T modalities only and full modalities*

### S8 Relationship between segmentation quality and image quality

In order to discover the difference between multi-modality and single-modality models, the volume of each subregion in each participant was calculated and was compared between models. Outliers (subregions with low consistency) were identified and analyzed to determine how they relate to image quality.

Figure S12 shows the subregion volume comparisons between these two models. The blue dots representing high quality ROIs (quality rating > 5) in each subplot of Figure S12 are distributed largely along the diagonal line. This demonstrates the consistency of segmentation results between single-modality and multi-modality models with ModAug in high-quality images. However, the red dots, which represent poor-quality ROIs, are not always distributed along the diagonal line in each subplot in Figure S12. Most outliers fall below the diagonal line, indicating that the segmentation results of the single-modality model produce smaller subregions than those of the multi-modality model. As can be seen from the examples in Figure S12, these outliers were poor quality images and segmentations. The single-modality model had severe under-segmentation, while the segmentation results from multi-modality model had typical segmentation size and morphology.

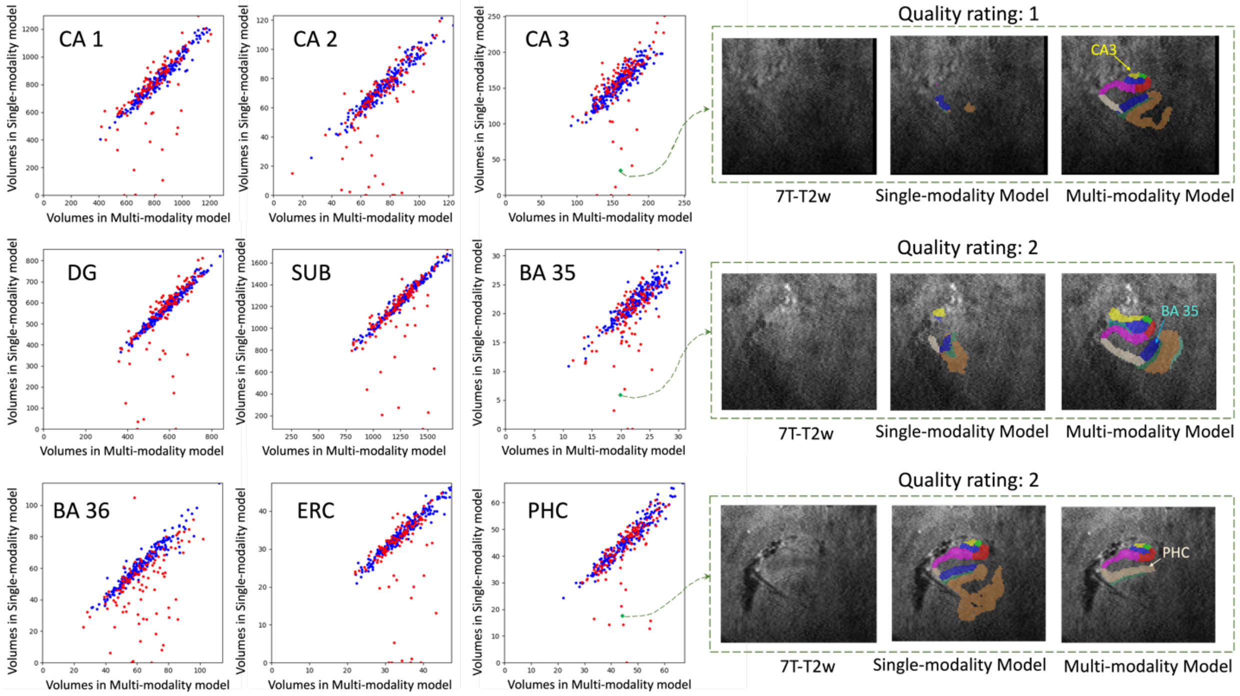


*Figure S12 The volume comparison for different subregions on the left side.* ***X-axis****: subregion volume by the multi-modality model with ModAug.* ***Y-axis****: subregion volume by the single-modality model. Blue dots: participants with good image quality. Red dots: participants with poor image quality.*

The relationship between image quality and segmentation quality was also explored using qualitative assessment results. Table S2 shows the segmentation quality of the two models in relation to image quality. The multi-modality model demonstrated greater robustness across various image qualities, which validated the visualization results presented in Figure S12. This finding further confirms that poor image quality was the reason why certain segmentations produced by the single-modality model should be excluded from downstream tasks.

*Table S2 Relationship between image quality and segmentation quality*

| | Single-modality | | | Multi-modality | | |
|---|---|---|---|---|---|---|
| **Seg. Quality** / **Image Quality** | Good | Uncertain | Bad | Good | Uncertain | Bad |
| **Low Quality** | 49 | 45 | 36 | 103 | 26 | 1 |
| **High Quality** | 189 | 25 | 0 | 197 | 17 | 0 |

### S9 Generalization performance on new datasets

We applied this pipeline on a new dataset that was just released in early 2025 (Chu et al., 2025), which contains 7T and 3T paired brain MRI from 20 subjects. Four main aspects of this dataset are different with our dataset:

- ***The 7T scanner is different***. Our dataset used 7T Siemens Terra scanner and the new dataset used 7T Siemens MAGNETOM scanner.
- ***The 7T-T1w acquisition protocol is different***. Our dataset used MP2RAGE with two inversion images and the new dataset used the standard T1w MRI.
- ***3T-T2w image resolution is different***. Although these two datasets used the same 3T scanner, the 3T-T2w images have different resolutions and coverages. The resolution of 3T-T2w image in our dataset is $0.4 \times 0.4 \times 1.2$ and it covers only partial region of brain; the resolution of 3T-T2w image in the new dataset is $0.86 \times 0.86 \times 1.88$ and it covers most of the brain region.
- ***The age of the participants is different***. Our test set has a mean age of 63 years (standard deviation=16.7, range = 23-94). The new dataset has a mean age of 23.35 years (standard deviation=1.42, range = 21-26).

We collected intermediate and final results of all steps in the pipeline when it was used for new dataset. Figure S13 is the example of these results on subject 01 in the new dataset. Since our model was trained with MP2RAGE 7T-T1w protocol, and the new dataset has standard T1w protocol, the modalities of 7T-T1w inv1 and inv2 were regarded as missing in the inference.

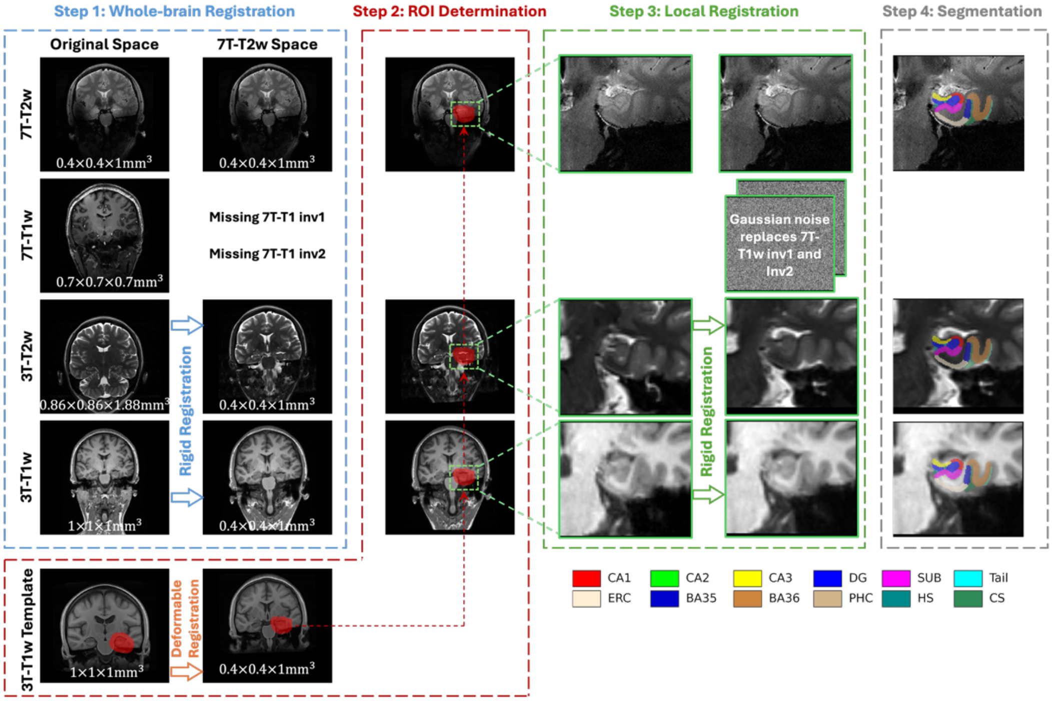


*Figure S13 Segmentation intermediate results of subject 01 in new dataset*

More examples are shown in Figure S14. We show one coronal view segmentation for the first five subjects and 3D rendering for the second five subjects. They all look good. These results can prove the successful inference of our pipeline on a new dataset. All segmentation results are available at https://doi.org/10.5061/dryad.0zpc8676p.

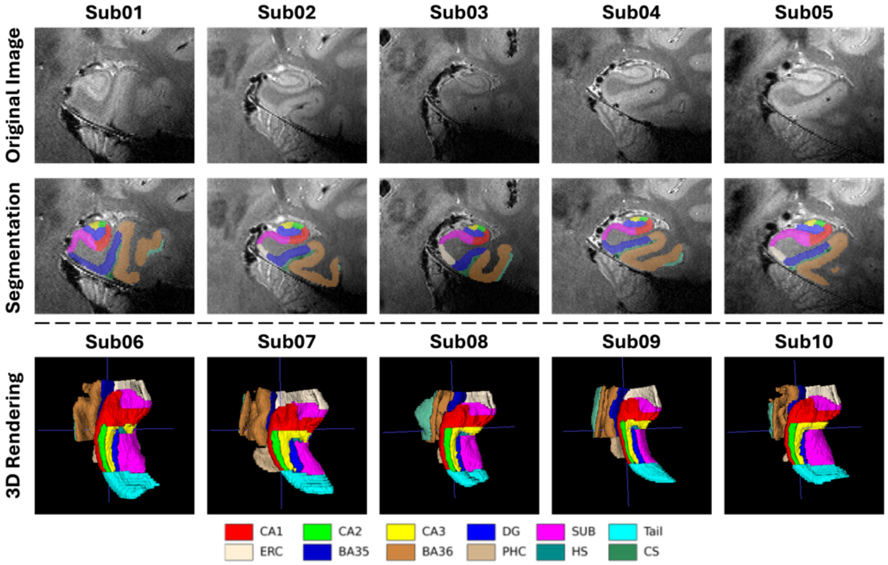


*Figure S14 Visualization segmentation in the new dataset*

### S10 Other modality augmentation methods

The purpose of modality augmentation is to force the model to extract consistent structural information from all available modalities, enabling the model to adapt to situations where some modalities are of low quality or missing during inference. There are various methods for modality augmentation. In addition to the random noise mentioned in the main text, we also compared the results of modality augmentation using images with all zeros and images from other participants.

The method of using all-zero images for modality augmentation is similar to that of using random noise. We randomly replace 1 to 4 modalities with all-zero images, and the remaining steps are the same.

For methods that use images from other participants for modality augmentation, we first need to establish a set of candidate images. We searched our dataset for participants who met the following two inclusion criteria:

- The participants have both 7T and 3T magnetic resonance images.
- The participants were not included in the training set or test set in previous experiments.

A total of 14 7T-3T paired images from 13 participants were selected. Their left and right MTL ROIs were extracted according to the ROI extraction process described in this paper, resulting in 28 candidate images. During the segmentation model training process, each modality to be replaced was randomly replaced by the corresponding modality from the 28 candidates. These new images were linearly resized to the training set image size and normalized using z-score normalization to match the mean and standard deviation of the replaced images. The specific workflow is shown in Figure S15 below.

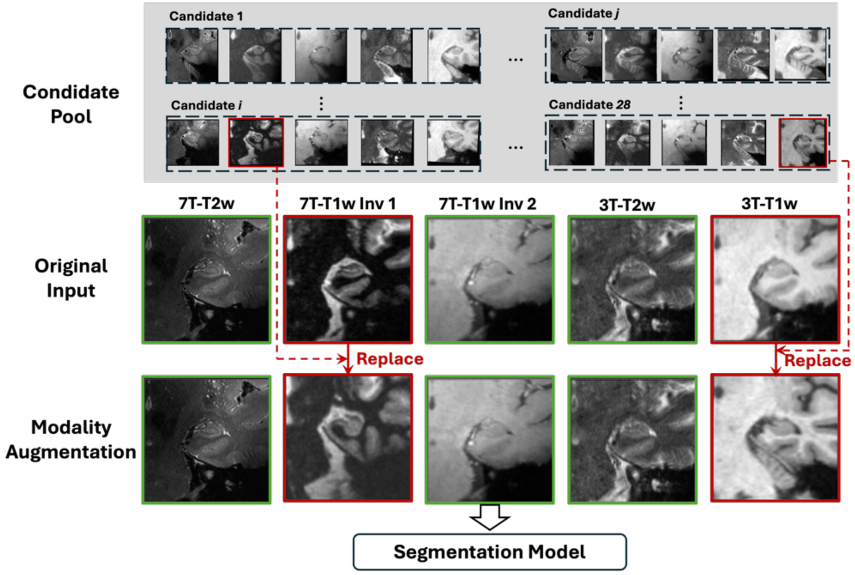


*Figure S15 The example of modality augmentation using other participants' images to replace certain modalities in training set*

We compared these three different modality augmentation methods' performance using Dice scores in five-fold cross-validation on the training set, the visualization results inferred from low-quality 7T-T2w samples in the test set, and the visualization results inferred from a third-party independent test set with missing 7T-T1w modality.

*Dice scores in five-fold cross-validation on the training set.* As shown in Table S3, when augmentation is performed using Gaussian noise, all-zero images, and images from other participants, the Dice scores for segmentation across different subregions are relatively consistent. Within each subregion, the differences between models are only in the third decimal place.This shows that during training, the disturbance information introduced by augmentation did not negatively affect the model's learning ability. Furthermore, the test data shows that these augmentation methods enabled the model to obtain stable outputs even when the main modality quality was low and modality was missing, achieving our goal of using augmentation to train the model to extract useful information from more modalities.

*Table S3 Dice scores of models with different augmentation methods in 5-fold cross-validation on the training set*

| Augmentation Method | Gaussian Noise | Zero Images | Other Images |
|---|---|---|---|
| **CA1** | 0.817 | 0.819 | 0.815 |
| **CA2** | 0.741 | 0.746 | 0.738 |
| **DG** | 0.866 | 0.869 | 0.862 |
| **CA3** | 0.724 | 0.728 | 0.720 |
| **Tail** | 0.861 | 0.860 | 0.858 |
| **SUB** | 0.862 | 0.865 | 0.861 |
| **ERC** | 0.855 | 0.856 | 0.851 |
| **BA35** | 0.751 | 0.752 | 0.751 |
| **BA36** | 0.825 | 0.827 | 0.826 |
| **PHC** | 0.822 | 0.821 | 0.821 |

*Segmentation visualization in low-quality test cases.* We also compared these modality augmentation methods in the test set with participants with low quality in the primary modality (7T-T2w). Figure S16 below shows three examples. It can be observed that the three modality augmentation methods achieve stable segmentation results under low-quality 7T-T2w images and exhibit consistency among themselves.

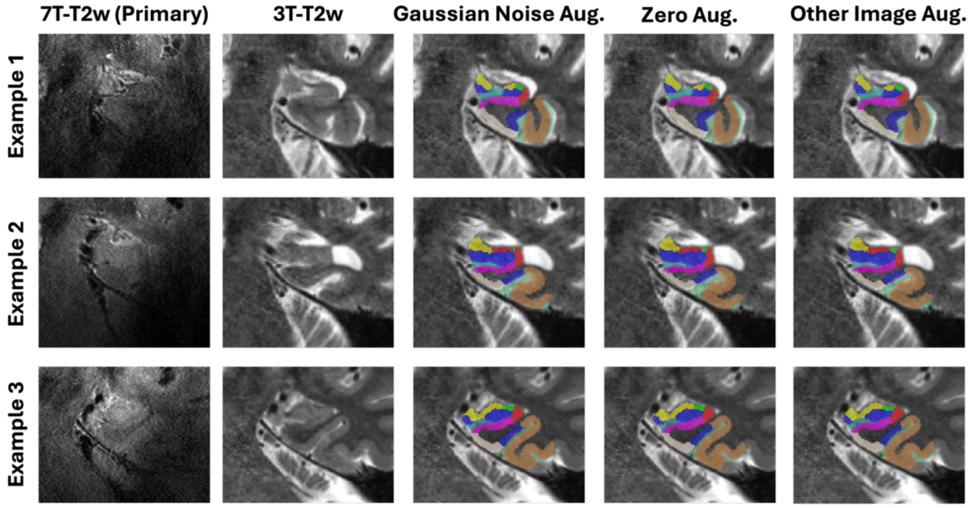


*Figure S16 Examples of segmentation results obtained using models trained with different modality augmentation methods in low-quality 7T-T2w scenarios. Due to the poor quality of the primary modality, the segmentation results are displayed in the 3T-T2w modality.*

*Segmentation visualization with missing modality in independent third-party test set*. Since third party dataset (Chu et al., 2025) uses a single 7T-T1w modality, rather than the 7T-T1w inv1 and 7T-T1w inv2 modalities used to train our model, we treat this dataset as missing the 7T-T1w inv1 and 7T-T1w inv2 modalities. In this case, we tested the performance of segmentation models trained under three modality augmentation strategies. Specifically, during inference, the 7T-T1w inv1 and 7T-T1w inv2 were replaced by each model's augmentation images. For example, for models that used images from other participants for augmentation, the missing modalities were replaced by images from other participants during inference as well. In this scenario, images of 7T-T2w, 3T-T2w, and 3T-T1w are all available, and we expect the model to identify consistency across modalities to complete segmentation. The segmentation examples are shown in Figure S17.

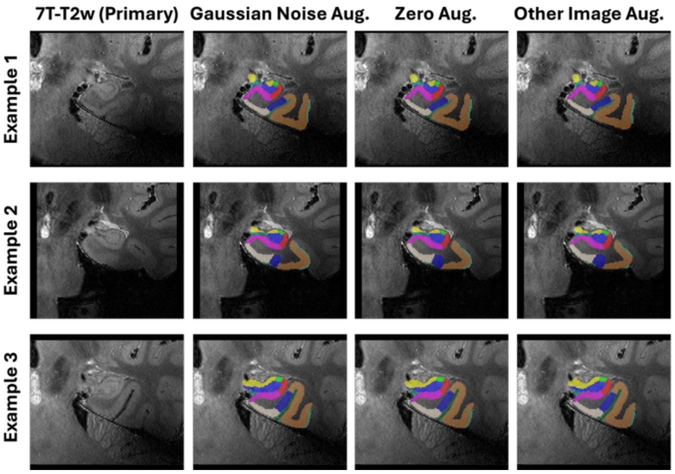


*Figure S17 Examples of segmentation inference from models trained under different modality augmentation strategies when 7T-T1w inv1 and 7T-T1w inv2 are missing.*

After comparison, all three modality augmentation methods achieved our goal, and the models trained using these methods were able to perform stable segmentation of the MTL subregions even when the main modality quality was poor and some modalities were missing.
